# Companion to: A molecular basis for milk allergen immune recognition in eosinophilic esophagitis

**DOI:** 10.1101/2024.11.13.623408

**Authors:** Julianna Dilollo, Alex Hu, Huiqi Qu, Karina E. Canziani, Rachel L. Clement, Sam J. McCright, Wayne G. Shreffler, Hakon Hakonarson, Jonathan M. Spergel, Karen Cerosaletti, David A. Hill

## Abstract

**Background:** Eosinophilic esophagitis (EoE) is a chronic, type 2 inflammatory disease that is increasing in incidence and has substantial morbidity. Despite being clinically defined as a food allergy, the molecular details of food antigen presentation and recognition by the immune system are unknown.

**Objective:** The objective of this study was to identify and characterize the molecular basis of milk antigen presentation and T cell recognition in a patient with EoE milk allergy.

**Methods:** Milk-expanded TCR clonotypes were identified using *ex vivo* stimulation followed by single cell RNA with linked, paired *TRA* and *TRB* sequencing. HLA restriction and antigen specificity of TCRs were identified using a combination of lentiviral expression, HLA sequencing, antibody blockade, and a peptide library screen.

**Results:** We isolated a β-casein specific TCR clonotype (eoeTCR-4) and determined its HLA restriction (HLA-DRB1*07:01) and cognate antigen (β-casein AA 59-78). EoeTCR-4 was not detected among a larger group of subjects and is likely private to EoE Subject 1.

**Conclusion:** In conjunction with the parent manuscript, this companion article provides the first molecular identity of food antigen presentation and immune recognition in EoE.

**SUMMARY:** This manuscript is a companion to the parent manuscript “A molecular basis for milk allergen immune recognition in eosinophilic esophagitis”. It provides additional data and detailed methods not provided in the parent manuscript. It is designed to supplement and support the findings and conclusions of the parent manuscript.

## INTRODUCTION

First described in 1978,(1) eosinophilic esophagitis (EoE) is a chronic inflammatory disease and one of the leading causes of dysphagia, odynophagia, and esophageal food impaction.(2,3) Since the early 1980s,(4) EoE has been managed as a clinical food allergy through the use of empiric food elimination diets.(5) This practice was supported by investigations which revealed that EoE has a strong predominance of type 2 (T2) allergic inflammation.(6,7) Despite this, immunologic evidence of food antigen recognition by the immune system in EoE was initially elusive, confounding clinical practice and impairing diagnostic and therapeutic advances.

In the early 2000s, the first data emerged for a central role for both CD4+ T helper type 2 (TH2) cells(8) and antigen-specific immune responses(9) in animal models of EoE. More recently, translational studies have identified food allergen-activated T cells in the circulation of patients with EoE, but not in healthy controls.(10–12) In addition, single cell RNA sequencing (scRNA-seq) of patient biopsies has elucidated the effector phenotype(13) and identified clonal expansion(14,15) of T cells. Despite these findings, alternative mechanistic explanations for EoE have been proposed including an epithelial cell–intrinsic alteration in TGFβ signaling that is independent of T cells.(16) As such, there is an active debate in the field as to the immunopathologic etiology of EoE.

Knowledge of the molecular basis of antigen presentation and recognition has clinical application in the diagnosis and management of inflammatory diseases. For example, the identification of specific autoantigens presented by MHC class II molecules to CD4+ T cells, and utilization of medications that interfere with T cell co-stimulation during antigen presentation, has revolutionized the diagnosis and management of rheumatoid arthritis.(17,18) In the case of IgE-mediated anaphylactic food allergy, knowledge of the specific antigenic components has increased diagnostic accuracy, allowed development of personalized anaphylaxis risk assessments for patients, and holds promise for targeted immunotherapy.(19,20)

We hypothesized that if EoE is a T cell-mediated, food-antigen driven disease, we should be able to identify the HLA allele and TCR clonotypes that mediate food antigen presentation and recognition by T cells. Further, we expect that in addition to expanding our understanding of disease etiology, identification of these molecules will have imminent clinical applicability to the development of novel antigen-specific diagnostics, therapeutics, and screening approaches. This manuscript provides additional data and detailed methods not provided in its parent manuscript. It is designed to supplement and support the findings and conclusions of its parent manuscript.

## DETAILED METHODS

### Study design and oversight

The objective of this study was to identify and characterize the molecular basis of milk antigen presentation and recognition in a patient with EoE milk allergy. This study complied with research protocols approved by the Institutional Review Board (IRB) for Human Subject Research at the Children’s Hospital of Philadelphia (IRB protocols 18-015524, DAH & 16-013278, HH), the Partners HealthCare at Massachusetts General Hospital (IRB protocol 2011P001159, WS), or the Human Immunology Core at the University of Pennsylvania. Informed consent and subject assent, where applicable, were obtained for all subjects.

### Study participants and sample acquisition

A subject with biopsy-confirmed, clinically diagnosed EoE milk allergy was identified and recruited for TCR and scRNA sequencing at the Children’s Hospital of Philadelphia. A board-certified pediatric allergist reviewed the subject chart prior to recruitment. EoE was defined in accordance with international diagnostic criteria throughout the manuscript.(21) Specifically, the subject was determined to have EoE milk allergy due to a normal upper endoscopy and esophageal biopsy (0-14 eosinophils/hpf) while avoiding milk only, and an upper endoscopy and esophageal biopsy displaying esophageal eosinophilia (≥15 eos/hpf) having introduced only milk into their diet. A peripheral blood sample (10-20 mL) was obtained in sodium heparin tubes, transported at room temperature to the lab, and immediately processed for culture. For lentiviral transfections, peripheral blood was obtained from healthy donors (Controls 1 and 2) and similarly processed. Additional control PBMCs were obtained from an apheresis donor via the Human Immunology Core of the University of Pennsylvania (Control 3).

### PBMC isolation, stimulation, and blockade

Peripheral blood mononuclear cells (PBMCs) were isolated by Ficoll gradient, and the sample from EoE Subject 1 was divided. Cells destined for flow cytometric analysis were carboxyfluorescein succinimidyl ester (CFSE) labeled (Invitrogen) and cultured at 1 million cells/mL in OpTimizer SFM (Gibco) for 6 days in a 96-well round-bottom plate (Corning) in the absence or presence of Tetanus toxoid (0.625 μg per 200 k cells, Astarte Biologics) or a cocktail of five endotoxin-depleted, purified milk protein preparations (6.25 μg of each of α-lactalbumin, β-lactoglobulin, α-casein, β-casein, and κ-casein per 200 k cells, Millipore Sigma), as previously described.(12) Cells destined for sequencing from EoE subject 1 were cultured in parallel without CFSE labeling. PBMCs from controls were isolated by Ficoll gradient or by apheresis prior to sorting of T cells for transduction or differentiation of monocyte derived dendritic cells.

### Flow cytometric analyses of stimulated PBMCs

After 6 days of culture, CFSE labeled PBMCs were harvested and enumerated. Cells were treated with Live/Dead Fixable Blue (Invitrogen) for 15 minutes at 4°C in the dark, washed, and stained with anti-human CD8 (SK1, BioLegend, 4:100, BV510), CD3 (SK7, eBioscience, 3:100, APC-Cy7, or BD Biosciences, 5:100, BUV395), CD4 (OKT4, BioLegend, 4:100, BV605), CD19 (HIB19, BioLegend, 4:100, AF700, or BioLegend, 4:100, BV711), CD45RA (HI100, BD Biosciences, 4:100, V450, or BioLegend, 4:100, BV786), and CD45RO (UCHL1, BioLegend, 5:100, BV650) for 30 minutes at 4°C in the dark. Cells were washed, fixed (Invitrogen), and permeabilized (Invitrogen) prior to overnight staining for interleukin (IL)-4 (MP4-25D2, BioLegend, 3:100, AF647). Data were acquired on a core maintained LSR Fortessa (BD) within 24 hours of harvest. The cytometer was compensated using OneComp eBeads (Invitrogen), unstained cells, CFSE stained cells, and Live/Dead Fixable Blue stained dead cells, as appropriate. 1 million events were acquired for each arm. Data were analyzed using FlowJo v8.1 software (Becton Dickinson). Memory CD4+ T cells were gated as forward and side scatter low (lymphocytes), Live/Dead negative, CD8−, CD19−, CD3+, CD4+, CD45RA−, CD45RO+.

### scRNA/TCR-Seq library preparation

After 6 days of culture, PBMCs from EoE Subject 1 were harvested and enumerated. The fraction destined for scRNA/TCR-seq analysis were subjected to a bead purification for CD4 and CD45RO (StemCell Technologies; REF 19157). CD4+/CD45RO+ cells from each experimental arm (unstimulated, milk stimulated) were loaded onto separate channels of the 10X Chromium Controller (10X Genomics), according to the manufacturer’s protocol, with a target capture of 10,000 cells per channel. Sequencing libraries were generated using the NextGEM Single Cell 5’ Kit v1.1 and the Chromium Single Cell Human TCR Amplification Kit. Gene expression and feature barcoding libraries were pooled at a ratio of 4:1. Sequencing of pooled libraries was carried out on a NovaSeq 6000 system (Illumina) using one NovaSeq 6000 S1 Reagent Kit v1.5 (100 cycles) with target depth of 25,000 raw reads per cell.

### scRNA/TCR-Seq Analyses

Sequencing data was quantified using 10x Genomics Cell Ranger 6.1.1(22) and aligned to the GRCh38 reference genome. Cell calling was performed using Cell Ranger’s default parameters. Genes detected in fewer than 10% of the cells were removed, and cells that contained fewer than 500 read counts, expression of fewer than 250 genes, or contained more than 25% mitochondrial reads were removed. Counts were normalized using the deconvolution method in the R package scran.(23) TCR sequence assembly, annotation, and clonotyping was performed using the cellranger VDJ pipeline with default parameters. Clonotypes were defined as a group of cells of which alpha and beta Complementarity-determining region 3 (CDR3) regions were sufficiently similar to be considered part of the same lineage (plus cells that only have an alpha or a beta chain), according to the Cell Ranger algorithm. Cells that shared a clonotype with at least one other cell were considered expanded. TCR sequence diversity was computed using the R package Immunarch(24) after downsampling to the same number of cells in milk-stimulated vs. unstimulated cultures. Dimensionality reduction and clustering analysis were performed in R using the Monocle3 package.(25–31) Differential expression analysis between clusters and between other experimental groups was also performed using the Monocle3 package. The peTH2 signature for each cell was defined as the mean z-score of log-transformed normalized expression of the following 22 genes: *IL4, IL5, IL13, PTGDR2, PLA2G16, PTGS2, HPGDS, PPARG, ACADVL, ACSL4, SLC27A2, LPCAT2, DGKE, GK, CHDH, ALOX5AP, GPR15, ICAM2, GATA3, IL17RB, FFAR3, IL1RL1.*(14) Differential expression analysis between cells expressing eoeTCR-4 and cells expressing milk-insensitive TCRs was performed using the LIMMA R package (doi:10.1093/nar/gkv007). Enrichment analysis on the Hallmark gene sets (<u>10.1016/j.cels.2015.12.004</u>) between clusters was performed using the *roast* (<u>10.1093/bioinformatics/btq401</u>) approach in the LIMMA R package (doi:10.1093/nar/gkv007). Enrichment was based on a model containing all clusters, using separate comparisons for each cluster vs. the mean gene expression in the remaining clusters.

### Lentiviral vector generation

Plasmids were synthesized (GenScript) encoding the rearranged *TRAV* and *TRBV* sequences cloned upstream of the murine *Trac* and *Trbc* constant regions in the lentiviral plasmid, pRLL-MND-GFP (Addgene plasmid #36247), replacing the GFP gene.(32) Insert sequences were synthesized and cloned using engineered Notl and BstZ17I restriction enzyme sites. Correct sequence and orientation were confirmed via Sanger sequencing, gel electrophoresis, and UV spec. The murine Trbc region was utilized for flow-cytometric detection. The CHOP Research Vector Core amplified all plasmids and packaged all vectors. Vectors were manufactured at the pilot scale with a third-generation packaging system and concentrated by centrifugation. Viral titers were assessed in HEK293 cells using a duplexed droplet digital PCR (ddPCR) with the viral packaging signal psi (Ψ) normalized to a housekeeping gene (RPP30), yielding the reported titers in transduction units (TU). Direct inoculation of bioburden plates was used to evaluate sterility.

### Lentiviral T cell transduction

PBMCs from control subjects were stained with Live/Dead Fixable Aqua (Invitrogen), CD8 (BV510), CD19 (BV711), CD3 (APC-Cy7), CD4 (BV605), CD11c (3.9, Biolegend, 5:100, BV650), CD45RA (BV786), and CD25 (BC96, BioLegend, 5:100, PE). Non-regulatory CD4+ T cells were sort purified with a FACSAria Fusion flow cytometer (BD) using the following strategy: Live/Dead Fixable Aqua/CD8 negative, CD19-, CD11c-, CD3+, CD4+, CD25-. The resulting population was majority positive for the naïve marker CD45RA. T cells were suspended at 1 million cells per mL in complete OpTimizer SFM supplemented with rhIL-2 (50 ng/mL, PeproTech) and protamine sulfate (10 µg/mL, MP Biomedicals), and stimulated with anti-CD3/28 beads (5 µL per 200 k cells, Gibco). The Multiplicity of Infection (MOI) was determined for each TCR vector by titration. The optimal MOI for all vectors was 20 µL per 200 k cells, resulting in 50-75% transduction efficiency, depending on the clone. rLV was added to culture medium at the same time as anti-CD3/28 beads and peak transduction was achieved within 48 hours. Transduced cells were expanded for 10 days, subcultured with fresh media containing rhIL-2 every 2-3 days, after which beads were removed by magnetic separation and cells were enumerated. Cells were rested in the absence of beads and rhIL-2 for 48 hours, then frozen at 10 million cells per mL of freeze medium (Fetal Bovine Serum, 10% Dimethylsulfoxide).

### Generation of monocyte-derived dendritic cells

Control PBMCs were suspended at 3 million cells per mL of OpTimizer SFM supplemented with 10% autologous serum, GM-CSF (55 ng/mL, R&D Systems), and rhIL-4 (0.2 µg/mL, PeproTech), and plated in a 12 well flat-bottom plate. After 48 hours, nonadherent cells were discarded with the culture medium and fresh media with cytokines was added to the monolayer of adherent cells. After 3 days of culture, media was discarded again, and fresh media with cytokines was added. After 24 hours, maturation cytokines IFNy (5 ng/mL, R&D Systems) and CL075 (0.4 µg/mL, InvivoGen) were added directly to culture medium. After 24 hours of maturation, monocyte-derived dendritic cells (moDC) were harvested by vigorous pipetting, enumerated, and frozen at 1 million cells per mL of freeze medium.

### Stimulation and flow cytometric analyses of TCR-transduced T cells

Transduced T cells and autologous moDCs were rapidly thawed, resuspended in complete OpTimizer SFM, and enumerated. moDCs were adjusted to 1 million cells per mL of media and evenly distributed to sterile FACS tubes with caps. moDCs were left untreated or antigen loaded by treating with 100 µg/mL of Soy protein extract (Stallergenes Greer), 100 µg/mL α-casein, 100 µg/mL β-casein, 100 µg/mL κ-casein, 100 µg/mL β-lactoglobulin, a cocktail of equal parts of the 3 caseins, 2 µg/mL tetanus toxoid, 0.1 µg/mL of recombinant β-casein (KanPro Research, Inc, no post-translational modifications), or 0.1 µg/mL of recombinant β-casein peptides (Alan Scientific) for 2 hours at 37°C prior to coculture with transduced T cells. Concurrently, T cells were CFSE labeled, adjusted to 1 million cells per mL of OpTimizer, and plated at 170 k cells per well in a 96 well round bottom plate. Antigen loaded moDCs were washed, resuspended at 1 million cells per mL OpTimizer, and added to wells containing T cells at 30 k cells per well. Some T cells were cultured in the absence of moDCs and left unstimulated or stimulated with 5 µL/well of anti-CD3/28 beads as negative and positive controls, respectively. Cultures were incubated at 37°C for 3 days, then harvested and stained with Live/Dead Fixable Blue, CD19 (BV711), CD3 (BUV395), CD4 (BV605), CD8 (BV510), and anti-mouse TCR beta (H57-597, Invitrogen, 5:100, PE). Data were acquired with an LSR Fortessa, analyzed using FlowJo, and gated by Live/Dead Fixable Blue negative, CD19-, CD8-, CD3+, CD4+ based on FMO controls.

### Synthesis of rβ-casein

Bovine β-casein A2 was cloned and expressed by KanPro Research, Inc. using E. coli. Plasmids were amplified by Inverse PCR. Synthesized gBLOCK was inserted directly between the NdeI and BamHI, linearized by PCR/Restriction digestion, using the one step plasmid by Infusion Cloning. Plasmids harboring β-casein protein were transformed into E coli Bl21(DE3)pRARE. Protein was expressed in LB medium in the presence of ampicillin and chloramphenicol with 0.4 mM IPTG at 37°C for 3-5 hours. Cells were pelleted and disrupted by freeze/thaw and sonication. Whole cell, soluble, and insoluble fractions were obtained by centrifugation at 195000 rpm for 30 minutes. Fractions of cells were loaded on a 10-20% gradient Tris-Glycine SDS-PAGE gel and stained with Instant Blue. β-casein A2 was over-expressed in the form of insoluble fraction as inclusion bodies. The purity of the inclusion bodies was about 80%. Insoluble inclusion bodies of β-casein A2 were dissolved in 50 mM Tris-HCl (pH 8.0) and 6 M urea. The solubilized proteins were loaded on a 5 mL Q HP (GE Healthcare) equilibrated with the same buffer. The protein was eluted from the column with 0-100% gradient Buffer of 1 M NaCl, 50 mM Tris-HCl (pH 8.0) without urea.

Fractions of cells were loaded on a 10-20% gradient Tris-Glycine SDS-PAGE gel and stained with Instant Blue. β-casein A2 was over-expressed in the form of insoluble fraction, as inclusion bodies can be purified from Q column. The purity of the inclusion bodies was >95%. Recombinant β-casein A2 purified in 50 mM Tris-HCl (pH 8.0) and about 150 mM NaCl by the ion exchange chromatography were further purified with Superdex S75 size exclusion column equilibrated with 20 mM Tris-HCl (pH 8.0) and 150 mM NaCl. β-casein A2 purified from Q column can be eluted from S75 as a single peak. The purity of the inclusion bodies was >95%. Fractions of first peak eluted from 5 runs were pooled, flash frozen in liquid nitrogen, and stored at -80°C. >95% pure, recombinant bovine β-casein A2 fractions were delivered at 1.1-3.5 mg/mL in 20 mM Tris-HCl (pH 8.0) and 150 mM NaCl.

### Synthesis of β-casein peptides

Peptides spanning the length of bovine β-casein from amino acid residue 59-224, encompassing the HLA-DRB1*07:01 binding sites predicted by NetMHCIIpan-4.0, were generated via Solid-Phase Peptide Synthesis by Alan Scientific Inc. Purity of all delivered peptides was 95%, and no N or C terminal modifications were added. Lyophilized peptides were reconstituted in DPBS -Ca/-Mg at 1 µg/µL and stored at -80°C.

### HLA restrictions

To test the HLA restriction of eoe-TCR4, moDC were incubated with recombinant β-casein (rβ-casein) at 0.1 µg/mL following 30 minutes of preincubation with either anti-HLA-DR (100 µg/mL), anti-HLA-DP (100 µg/mL), anti-HLA-DQ (100 µg/mL), all 3 blocking antibodies, or isotype control to anti-HLA-DR (100 µg/mL). After 2 hours of incubation with antigen in the absence or presence of blocking antibodies, moDCs were washed and cocultured with CFSE labeled T cells transduced with eoeTCR-4. To test the restriction of rβ-casein presentation by EoE Subject 1 PBMCs, anti-HLA-DR (L243, BioLegend, 0.5 µg per 200 k cells) was added to an arm of the rβ-casein (6.25 μg per 200 k cells) culture condition.

### Antigen binding prediction

Binding sites were predicted for EoE Subject 1’s HLA-DRB1 alleles, 07:01 and 11:04, for each of the five major milk proteins (including both subunits of α-casein) using the Technical University of Denmark’s NetMHCIIpan-4.0 tool. NetMHCIIpan-4.0 is an HLA class II bind prediction server freely available online.(33) Milk protein sequences were submitted in FASTA format, peptide length was set to 15 amino acids, the threshold for a strong bind was set to <1% Rank, and the threshold for a weak bind was set to <5% Rank. Server output was displayed as a heatmap for each protein in order of predicted bind position.

### Mass spectrometry data acquisition

Each of the five Millipore Sigma purified bovine milk proteins, endotoxin depleted and adjusted to 5 µg/µL in DPBS -Ca/-Mg, were submitted to the CHOP Proteomics Core for compositional analysis by Mass Spectrometry. Proteins were solubilized and digested with the iST kit (PreOmics GmbH, Martinsried, Germany) per manufacturer’s protocol.(34) The resulting peptides were de-salted, dried by vacuum centrifugation and reconstituted in 0.1% TFA containing iRT peptides (Biognosys Schlieren, Switzerland). Peptides were analyzed on a Q Exactive HF mass spectrometer (ThermoFisher Scientific San Jose, CA) coupled with an Ultimate 3000 nano UPLC system and an EasySpray source using data dependent acquisition (DDA). Peptides were loaded onto an Acclaim PepMap 100 75 μm x 2 cm trap column (Thermo) at 5 μL/min and separated by reverse phase (RP)-HPLC on a nanocapillary column, 75 μm id × 50 cm 2 μm PepMap RSLC C18 column (Thermo). Mobile phase A consisted of 0.1% formic acid and mobile phase B of 0.1% formic acid/acetonitrile. Peptides were eluted into the mass spectrometer at 300 nL/min with each RP-LC run comprising a 90-minute gradient from 3% B to 45% B. The mass spectrometer was set to repetitively scan m/z from 300 to 1400 (R = 240,000) followed by data-dependent MS/MS scans on the twenty most abundant ions, minimum AGC 1e4, dynamic exclusion with a repeat count of 1, repeat duration of 30 seconds, and resolution of 15000 ions. The AGC target value was 3e6 and 1e5 ions, for full and MSn scans, respectively. MSn injection time was 160 ms. Rejection of unassigned and 1+,6-8 charge states was set.

### Mass spectrometry system suitability, quality control, and data analysis

The suitability of Q Exactive HF instrument was monitored using QuiC software (Biognosys, Schlieren, Switzerland) for the analysis of the spiked-in iRT peptides. As a measure for quality control, standard E. coli protein digest was injected prior to and after injecting the sample set and collected the data in the Data Dependent Acquisition (DDA) mode. The collected DDA data were analyzed in MaxQuant,(35) and the output was subsequently visualized using the PTXQC(36) package to track the quality of the instrumentation. MS/MS raw files were searched against a bos taurus protein sequence database including isoforms from the Uniprot Knowledgebase (taxonomy:9913, 37501 entries) using MaxQuant version 1.6.14.0 with the following set parameters: Fixed modifications, Carbamidomethyl (C); Decoy mode, revert; MS/MS tolerance FTMS 20 ppm; false discovery rate for both peptides and proteins, 1.0; Minimum peptide Length, 7; Modifications included in protein quantification, Acetyl (Protein N-term), Oxidation (M); Peptides used for protein quantification, Razor and unique. Scaffold (version Scaffold_5.1.0, Proteome Software Inc., Portland, OR) was used to validate MS/MS based peptide and protein identifications. Peptide identifications were accepted if they could be established at greater than 95.0% probability by the Percolator posterior error probability calculation.(37) Protein identifications were accepted if they could be established at greater than 99.0% probability and contained at least 1 identified peptide. Protein probabilities were assigned by the Protein Prophet algorithm.(38) Proteins that contained similar peptides, and could not be differentiated based on MS/MS analysis alone, were grouped to satisfy the principles of parsimony. Proteins sharing significant peptide evidence were grouped into clusters. Total Spectrum Counts were exported from Scaffold and the 218 identified proteins across all five samples were grouped into categories: one of the expected milk proteins (α-casein subunits 1 or 2, β-casein, κ-casein, α-lactalbumin, or β-lactoglobulin), other bovine proteins, the mass spectrometry control protein, or non-bovine contaminants.

### HLA genotyping and enrichment analysis

EoE Subject 1 Control 1, and Control 2’s HLA types were determined by 11-loci Next Generation Sequencing at the American Red Cross HLA Laboratory in Philadelphia, PA. Control 3 was HLA typed by the same method at The Clinical Immunology and HLA Immunogenetics Laboratory at Hospital of the University of Pennsylvania.

### Characterization of CDR3 regions by high-throughput sequencing

Genomic DNA was extracted from whole blood using a standard protocol. Whole exome sequencing was performed at the Center for Applied Genomics of the Children’s Hospital of Philadelphia. Paired-end sequencing was performed on the Illumina NovaSeq 6000 platform (Illumina, San Diego, CA), using an S4 flowcell with run parameters of 101 x 10 x 10 x101 [Read 1 x Index 1 (i7) x Index 2 (i5) x Read 2]. All sequencing experiments adhered to quality control standards of depth of coverage, read quality, and error rates. The characterization of *TRA and TRB* CDR3 regions was performed using raw sequencing reads in FASTQ format with miTCR.(39) A threshold of a minimum quality score of 25 for Illumina sequencing reads was used to ensure that only high-quality reads are used for inferring TCRs.

### Comparison of CDR3 regions by scTCR-seq

We computed pairwise amino acid sequence similarity of 43,282 *TCRB* and 27,534 *TCRA* sequences from esophageal tissue and PBMC samples obtained from patients with EoE milk allergy. This was achieved by generating Levenshtein distance matrices of the CDR3 sequences using the Stringdist package in R.(40) The results were confirmed by computing weighted multi-CDR distances between TCRs using Tcrdist3, a Python3 package for TCR repertoire analysis, following the procedure described in Mayer-Blackwell et al., 2021.(41)

### Data and materials availability

Data are available in the main text, supplementary materials, or upon reasonable request.

## RESULTS

To begin our search for a food specific T cell, we collected peripheral blood samples from a subject with clinically confirmed EoE caused by exposure to cow’s milk (EoE milk allergy) (EoE Subject 1, **Fig. ED1a**). EoE Subject 1 had a “classic” presentation, having developed EoE in early childhood with multiple subsequent periods of milk removal and reintroduction with corresponding remission and reactivation of his EoE (**Fig. ED1b**). We then performed scRNA-seq with linked, paired *TRA* and *TRB* sequencing (scTCR-seq) to determine the activation states and TCR clonotypes of TH cells that expanded upon milk protein stimulation (**Fig. 1a**).

**Fig. 1.**
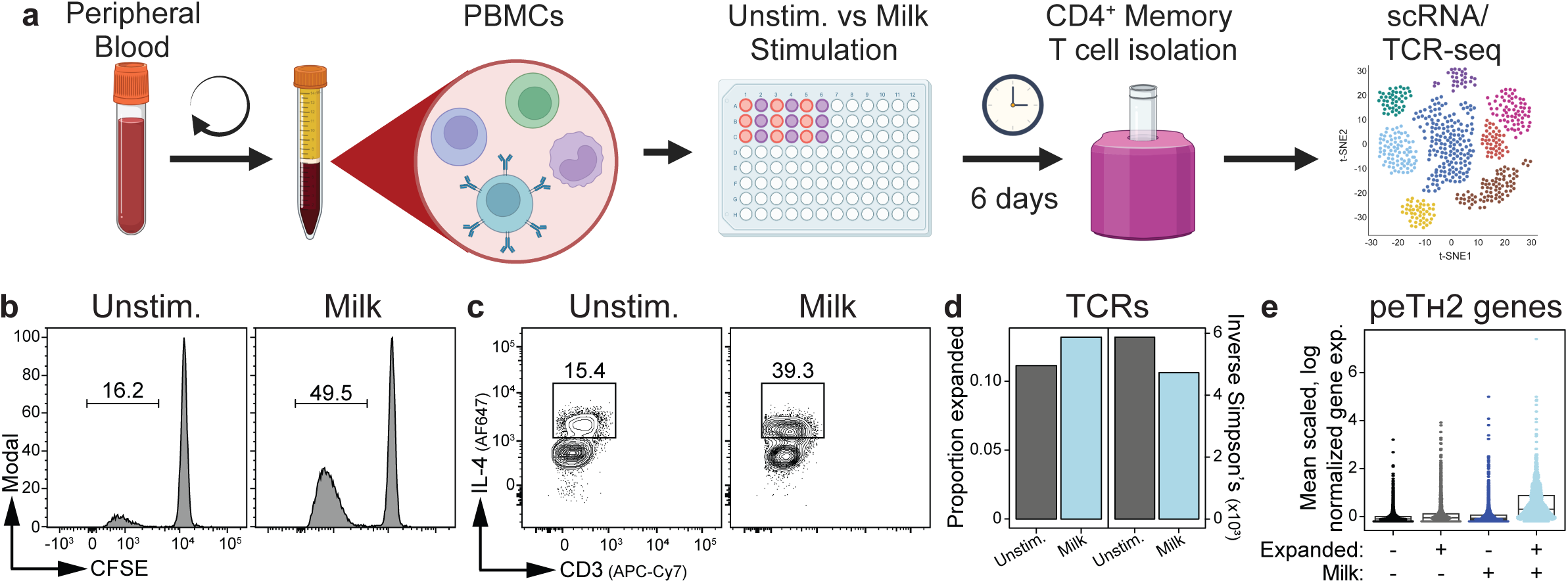
Identification of milk-expanded TCR clonotypes from a child with EoE milk allergy. (**a**) PBMC from EoE Subject 1 were unstimulated (Unstim.) or stimulated with a cocktail of α−casein1, α−casein2, β−casein, κ−casein, α−lactalbumin, and β−lactoglobulin (Milk) for 6 days. CD4^+^ memory T cells were bead-purified and subjected to single cell RNA sequencing with linked TCR sequencing. (**b**) Carboxyfluorescein succinimidyl ester (CFSE) fluorescence intensity of unstimulated or milk-stimulated T cells from EoE Subject 1. (**c**) Interleukin 4 (IL-4) expression by unstimulated or milk-stimulated T cells from EoE Subject 1. (**d**) TCR expansion and diversity (by inverse Simpson’s score) between unstimulated and milk-stimulated cultures from EoE Subject 1. Data represent the mean of 100 down-samples. (**e**) Distribution of the peTH2 gene signature per cell and by expansion state and stimulation condition in EoE Subject 1. Data shown as mean z-score of log-normalized expression. All experimental arms are significantly statistically different from one another (p < 10^-16^).

Milk protein stimulation of PBMCs from EoE Subject 1 caused a subset of CD4+ CD45RO+ memory TH cells to proliferate and produce IL-4 (**Fig. 1b,c** and **Fig. ED2**).(12) To quantitate clonal expansion, we downsampled to equivalent numbers of cells per condition and determined the proportion of cells expanded and their TCR diversity using the inverse Simpson’s diversity index. This analysis confirmed increased clonal expansion and reduced TCR diversity upon milk stimulation (**Fig. 1d**).

Previous scRNA-seq analysis of esophageal biopsies from milk allergic EoE patients identified clonally expanded pathogenic effector TH2 cells (peTH2).(14) We therefore examined expression of peTH2 genes in our datasets (**Fig. ED3**). As anticipated, clonally expanded cells from milk-stimulated cultures had the highest expression of peTH2 genes compared with other conditions (**Fig. 1e**). To maximize our likelihood of identifying a milk-specific TCR from a pathogenic cell, we concentrated our validation efforts on milk-expanded clonotypes that expressed the peTH2 gene signature. Among the 20 most expanded TCR pairs in milk-stimulated culture from EoE Subject 1 (**Extended Data Table 1**), we detected high expression of the peTH2 gene signature in several clonotypes (data in parent manuscript). Of these, we chose eoeTCR-2 (*TRA* CVVVGRTGGFKTIF), 4, 6, and 7 for functional testing based on their uniqueness from one-another, absence of predicted specificities, and the peTH2 gene signature of the related T cell clones.

We determined the *HLA-DRB1* genotype of EoE Subject 1 (07:01, 11:04) and found a control donor with a near *HLA-DRB1* match (07:01, 11:01) (Control 1, **Extended Data Table 2**). CD4+ TH cells and autologous monocyte-derived dendritic cells (moDCs) from Control 1 were used as a TCR expression and testing system to establish the specificity of TCR clonotypes of interest.

TCR clonotypes were transduced into sort purified CD25-CD4+ conventional (Tconv) cells from Control 1 PBMCs using lentiviral vectors in which the recombined *TRA* and *TRB* V-CDR3-J region was cloned upstream of the murine *Trac* and *Trbc* region gene segments (**Fig. 2a**).(32) We then tested transduced Tconv cells for proliferation when cultured in the presence of protein pulsed autologous moDCs. Tconv cells expressing eoeTCR-2 (*TRA* CVVVGRTGGFKTIF), 6, and 7 were not reactive to milk proteins (**Fig. ED4** and not shown). However, Tconv cells expressing eoeTCR-4 showed robust proliferation when cultured in the presence of moDCs pulsed with α, β, or κ-casein, but not β-lactoglobulin, soy protein extract, or tetanus toxoid (**Fig. 2b**). These results indicate that eoeTCR-4 is casein-specific.

**Fig. 2.**
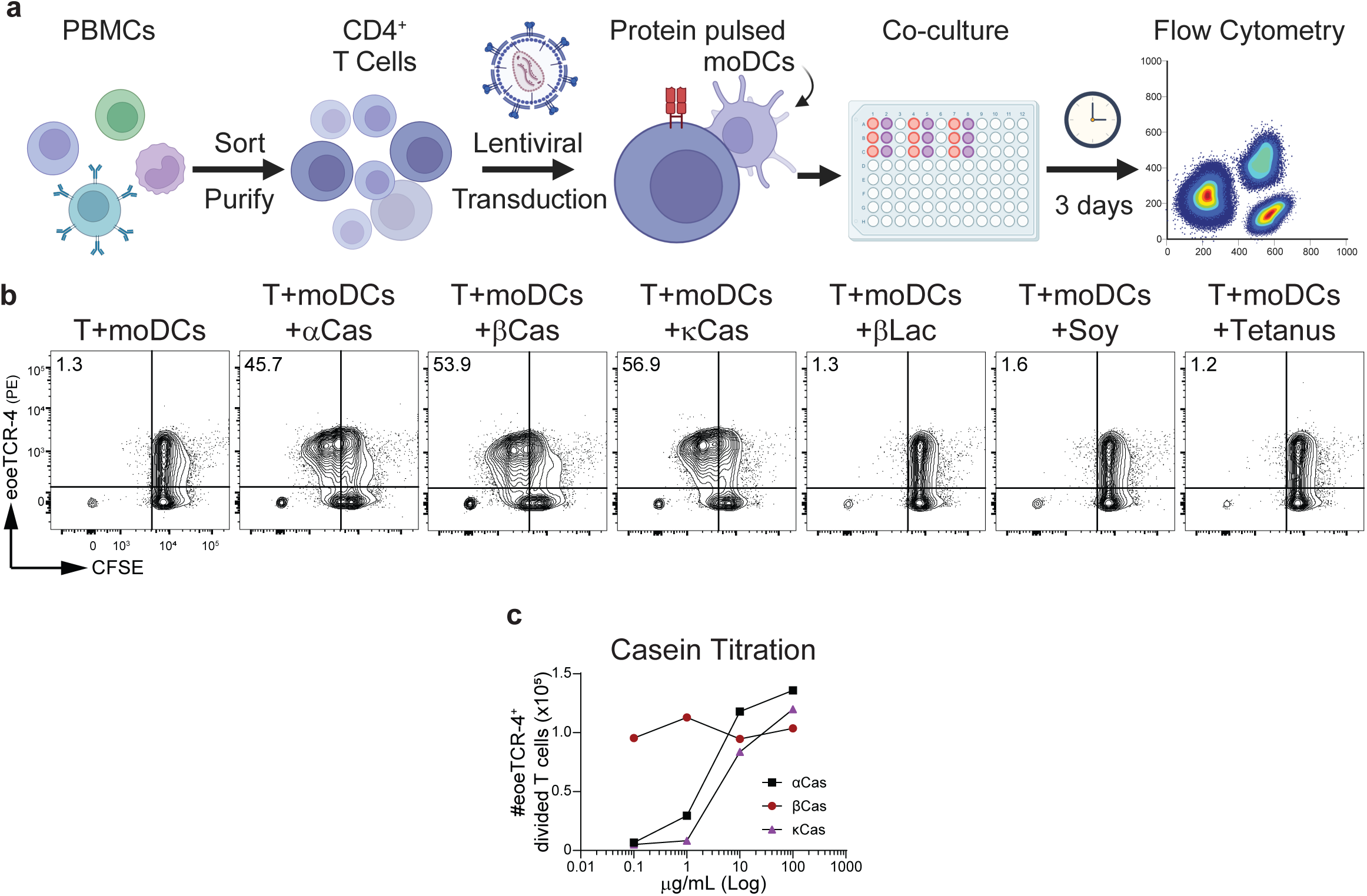
Identification of a β−casein-specific TCR clonotype from a child with EoE milk allergy. (**a**) CD4^+^ conventional T (T) cells were sort-purified from a HLA-DR-matched subject, transduced with lentiviral vectors encoding the TCR clonotypes of interest, and co-cultured with autologous, protein-pulsed monocyte-derived dendritic cells (moDCs) for three days prior to flow-cytometric evaluation. (**b**) Carboxyfluorescein succinimidyl ester (CFSE) fluorescence intensity of T cells transduced with eoeTCR-4 and cultured with moDCs alone or moDCs pulsed with α−casein subunit 1 and α−casein subunit 2 (αCas), β−casein (βCas), κ−casein (κCas), β−lactoglobulin (βLac), soy proteins (soy), or tetanus toxoid (Tetanus). (**c**) Cell division of T cells transduced with eoeTCR-4 and cultured with moDCs pulsed with α, β, or κCas. Data representative of 2 or more independent experiments.

We considered if the observed cross-reactivity of eoeTCR-4 to α, β, and κ-casein was due to epitope conservation across the proteins. However, machine learning predictions of potential binding sites for EoE Subject 1’s HLA-DRB1 alleles suggested that epitopes from κ-casein were unlikely to be presented by either DRB1*07:01 or 11:04 (**Fig. ED5a**). Alternatively, we considered the presence of cross-contamination of the caseins in the protein preparations we were using in culture. Indeed, all three casein products contained α-casein subunit 1 and 2, β-casein, and κ-casein, while the whey protein β-lactoglobulin was not contaminated with casein (**Fig. ED5b**). We next performed a limiting dilution of each of the casein products and found that β-casein caused proliferation of eoeTCR-4+ T cells at low concentrations while α and κ-casein did not (**Fig. 2c**). Together, these data identify β-casein as the most likely target of eoeTCR-4. To confirm this hypothesis, we obtained recombinant β-casein without post-translational modifications (rβ-casein) and compared stimulation of eoeTCR-4+ T cells with purified β-casein to rβ-casein. Similar responses from transduced cells to both native and rβ-casein established that β-casein contains the cognate antigen of eoeTCR-4 (data in parent manuscript).

We next sought to determine the HLA-DR allelic restriction for eoeTCR-4. To do so, we first confirmed HLA-DR restriction of eoeTCR-4 using rβ-casein and HLA class II blocking antibodies, showing that only anti-DR antibody blocked the response of eoeTCR-4 to rβ-casein (**Fig. 3a**). In addition, we confirmed that memory TH cells isolated from EoE Subject 1 were reactive to rβ-casein, and that the rβ-casein proliferative response of these TH cells was dependent on HLA-DR (**Fig. 3b**). Finally, we recruited three additional control subjects that shared either the HLA-DRB1*07:01 allele of EoE Subject 1 (Control 2), approximated the HLA-DRB1*11:04 allele of EoE Subject 1 and was positive for HLA-DRB3*02:02 (Control 3), or was positive for the HLA-DRB4*01:03 allele in the absence of HLA-DRB1*07:01 (Control 4) (**Extended Data Table 2**). Culture of eoeTCR-4 transduced Tconv cells with β-casein pulsed autologous moDCs from Control 2 (07:01, 13:01) resulted in T cell proliferation (**Fig. ED6a**), while culture of eoeTCR-4 transduced Tconv cells with β-casein pulsed autologous antigen presenting cells (APCs) from Control 3 (DRB1*11:03, DRB3*02:02) or Control 4 (DRB4*01:03) did not (**Fig. ED6b,c**). Together, these data indicate that eoeTCR-4 is restricted to HLA-DRB1*07:01.

**Fig. 3.**
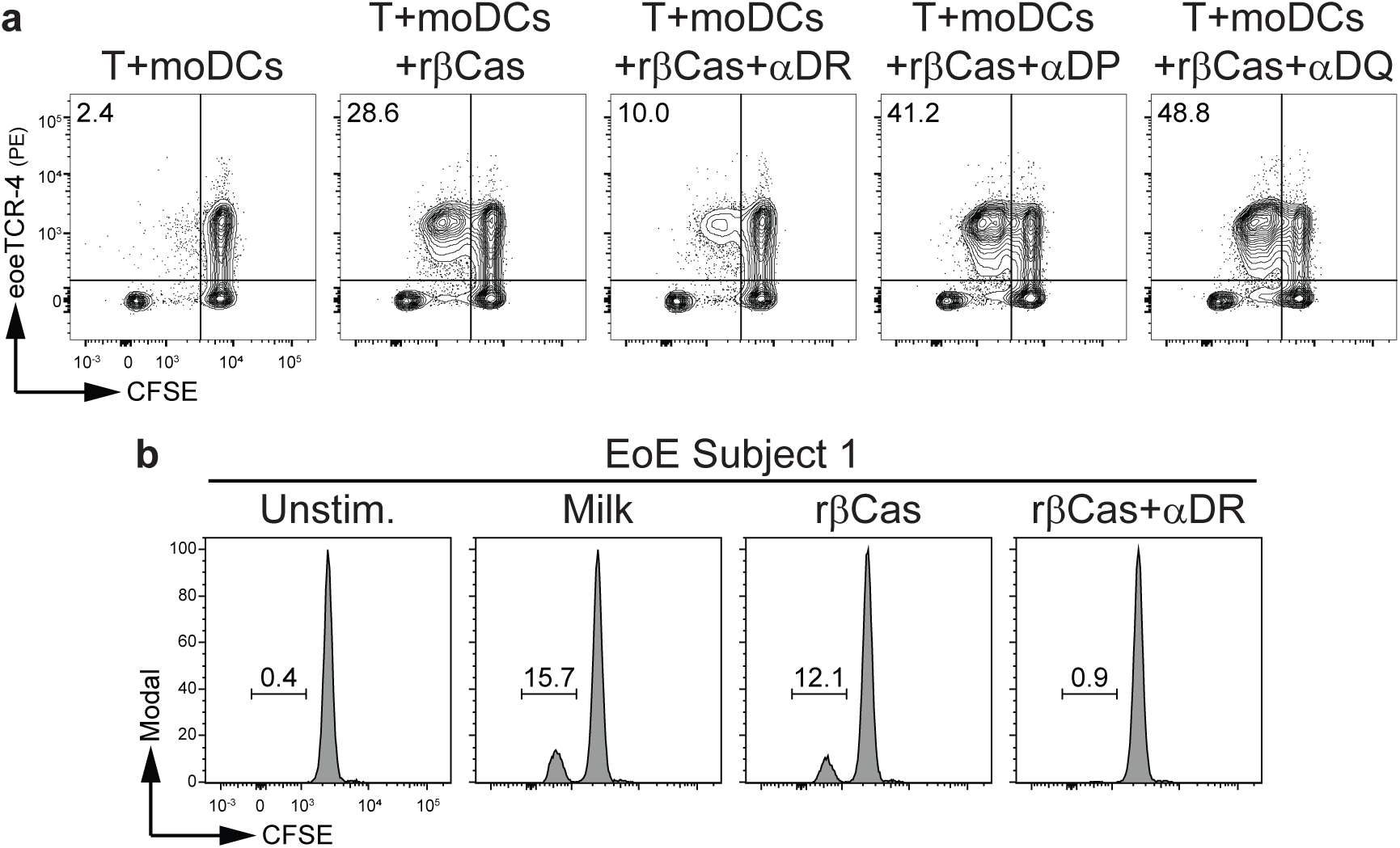
eoeTCR-4 is restricted to HLA-DR. (**a**) Carboxyfluorescein succinimidyl ester (CFSE) fluorescence intensity of CD4^+^ conventional T (T) cells transduced with eoeTCR-4 and cultured with autologous, protein-pulsed monocyte-derived dendritic cells (moDCs) alone or moDCs pulsed with recombinant βCas (rβCas) in the absence or presence of HLA-DR (αDR), -DP (αDP), or -DQ (αDQ) blocking antibodies. Data representative of 2 or more independent experiments. (**b**) CFSE fluorescence intensity of unstimulated, milk, or rβCas-stimulated T cells from EoE Subject 1 cultured in the absence or presence of αDR. Data representative of 1 experiment.

Having established that eoeTCR-4 is specific for β-casein and restricted to HLA-DRB1*07:01, we next sought to determine its cognate antigen. To do so, we contracted a third party to synthesize a peptide library that spanned the entirety of β-casein (**Fig. 4a**). Unfortunately, they were unable to express the first 58 amino acids of the protein. However, they were able to synthesize an overlapping peptide library of 9 peptides that spanned amino acids 59-224 and contained all three of the predicted β-casein binding sites (**Fig. 4a** and **Extended Data Table 3**). We then tested reactivity for each of these peptides with eoeTCR-4 using our lentiviral transduction system. We found that peptide 1, which contained the DRB1*07:01 predicted binding site 1, caused specific proliferation of eoeTCR-4 transduced T cells (**Fig. 4b**), while the remaining peptides did not (**Fig. 4b** and **Fig. ED7**). Together, these data identify β-casein peptide 1 as containing the cognate antigen for eoeTCR-4.

**Fig. 4.**
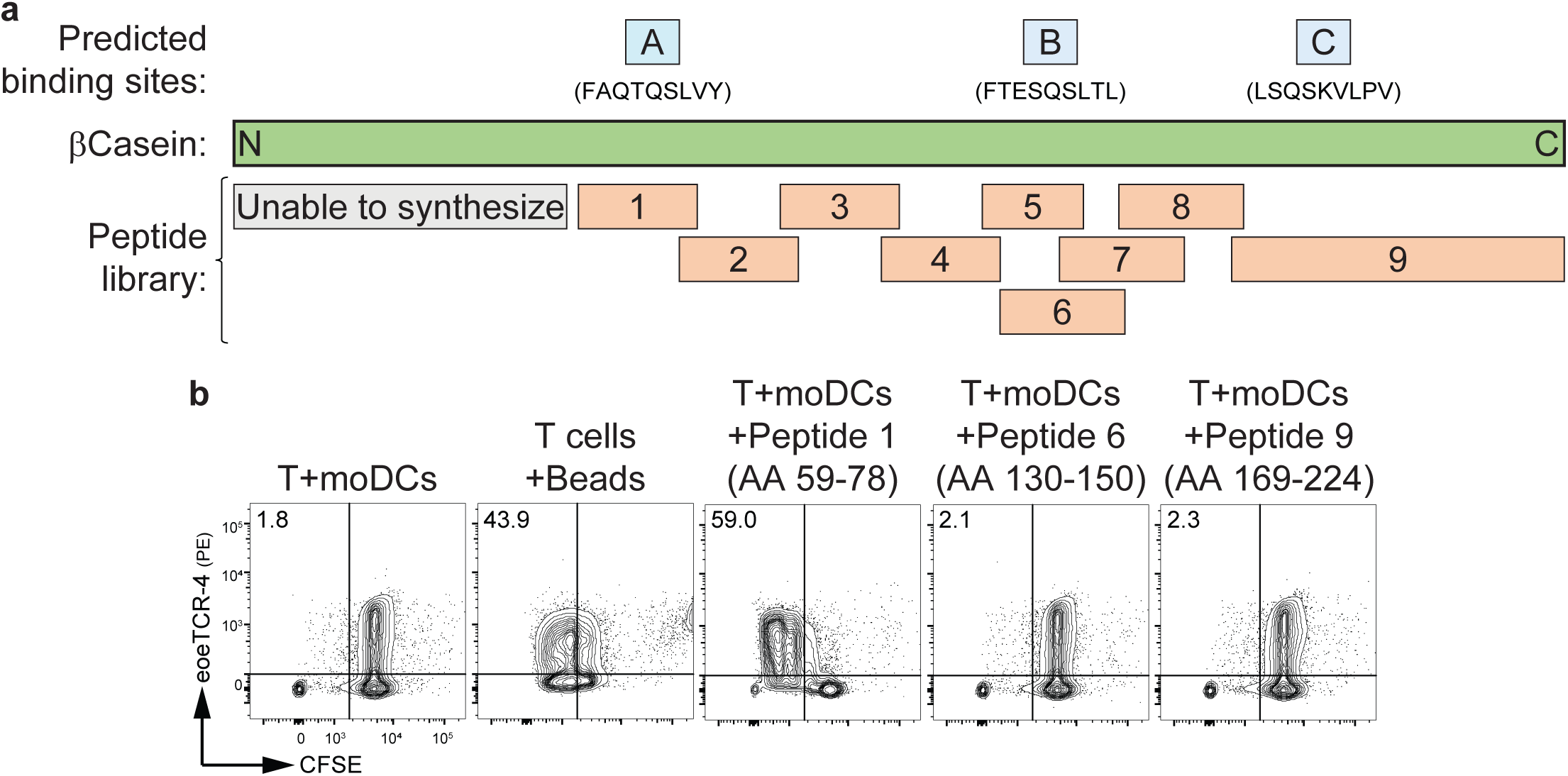
Identification of the cognate antigen for eoeTCR-4. (**a**) β−casein protein with predicted HLA-DR*07:01 binding sites (blue, as determined by NetMHCIIpan, A-C) and peptide library (red, 1-9). (**b**) Carboxyfluorescein succinimidyl ester (CFSE) fluorescence intensity of T cells transduced with eoeTCR-4 and cultured with moDCs alone, anti-CD3 anti-CD28 activating beads (beads), or moDCs pulsed with β−casein peptides 1, 6, or 9. Data representative of 2 or more independent experiments.

We next sought to determine the extent to which eoeTCR-4 was shared among patients with EoE milk allergy. First, we inferred *TRA* and *TRB* sequences from whole exome sequencing data of 132 unique patients with EoE milk allergy (**Extended Data Table 4**).(42–44) Among these patients, we did not observe an amino acid sequence match for either the *TRA* or *TRB* gene sequence of eoeTCR-4. Second, we examined a set of single cell TCR sequences from 21 patients, enriched for potential specificity (e.g., CD154+ following *in vitro* stimulation, expansion in tissue during active EoE, or expression of CD161+ CRTH2+ in circulation) from subjects with EoE milk allergy.(14) Using homology searching (Levenshtein distance, not shown; tcrdist3, **Fig. ED8**), we did not detect eoeTCR-4 or related sequences.

Finally, we sought to understand the extent to which there were unique transcriptional features of our milk-expanded clonotypes. We first performed unbiased clustering of CD4+ T cells from both unstimulated and milk stimulated cultures (data in parent manuscript and **Fig. ED9a**). We then overlayed clone groups based on their culture condition and expansion state, specifically noting the 20 most expanded clones in milk-stimulated culture as well as eoeTCR-4 clones (data in parent manuscript and **Fig. ED9b**). We noted that cluster 5 contained the highest proportion of the top milk-expanded clones, including eoeTCR-4 (**Fig. 5**). Cluster Hallmark gene set enrichment analysis revealed enrichment for interferon alpha and gamma response genes in cluster 5, 3, and 8, along with a generally pro-inflammatory transcriptome in the latter clusters (data in parent manuscript). In addition, we performed a transcriptional analysis of cells expressing eoeTCR-4, as compared with cells from other conditions and expansion states. We found that cells expressing eoeTCR-4 were most similar to other expanded cells from the milk condition (**Fig. ED10**). Finally, we performed a transcriptional analysis of cells expressing eoeTCR-4 compared to milk-expanded cells expressing TCR clonotypes that we established to be non-milk-specific (eoeTCR-6 and 7). Of note, cells expressing the eoeTCR-2 clonotype were not included in this analysis as we only tested one of the two expressed *TRA* chains. This final analysis revealed that cells expressing eoeTCR-4 had a unique transcriptome, with notably elevated expression of genes related to activation (*TMSB4X*, *S100A4*), cytotoxicity (*SRGN*), exhaustion (*CAPG*, *S100A10*), and regulation (*NDFIP1*) (data in parent manuscript).

**Fig. 5.**
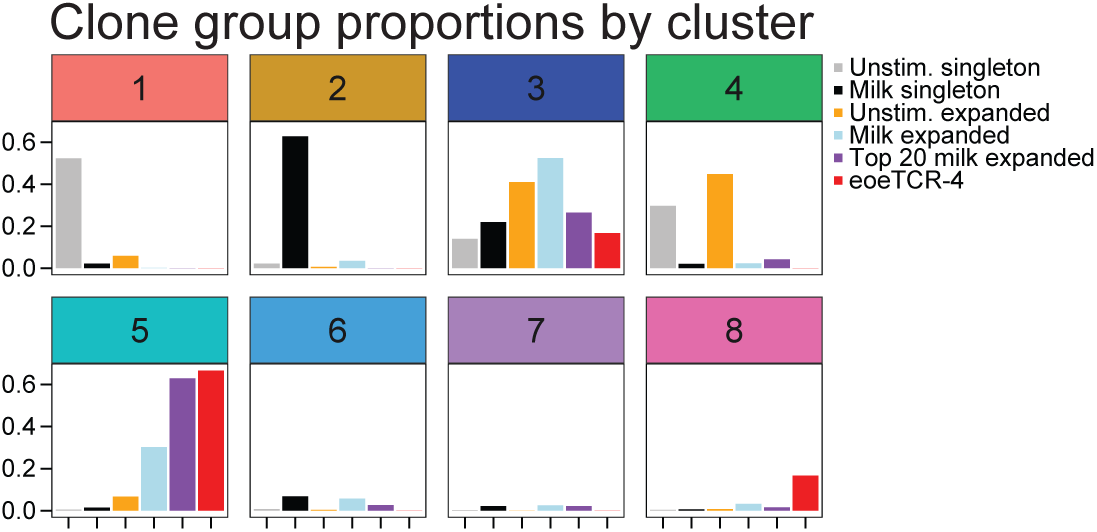
Milk-expanded T cell clonotypes are enriched for interferon response gene signatures. Clonotype group proportions in each CD4+ memory T cell cluster.

## DISCUSSION

The data provided herein describe the first molecular details of food allergen presentation and recognition by T cells in EoE, including the molecular identity of an EoE antigenic epitope (**Fig. 6**). Building from seminal studies of the EoE T cell repertoire from the past few years,(12–15) and taken in context with early studies indicating a very high response rate of EoE to elemental and milk elimination diets,(45–47) our findings support existing data indicating that a large majority of EoE is likely caused by food antigen-specific activation of T cells.

**Fig. 6.**
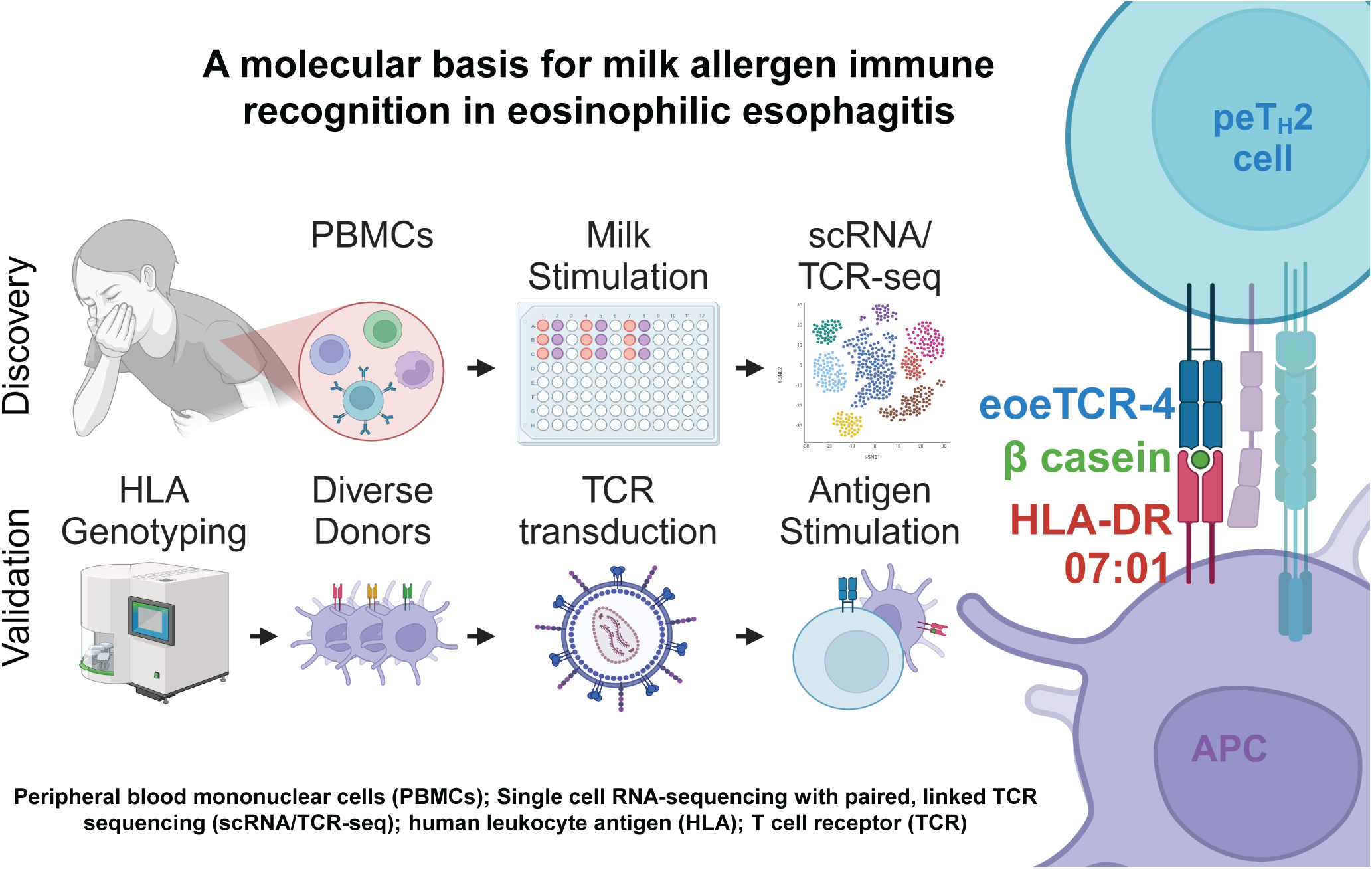
Graphical abstract. Created in BioRender. Hill, D. (2024) https://BioRender.com/d65u729.

Our findings support decades of clinical practice focused on food allergen identification and elimination in EoE. Perhaps most notably, we find that the peTH2 transcriptional signature was preserved in milk-expanded circulating memory T cells, including cells expressing eoeTCR-4. Our work also identifies novel features of the EoE T cell. For example, we find that circulating T cells that expanded upon milk stimulation were more likely to show transcriptional evidence of having responded to interferons. This finding is consistent with our recent observation that there is a conserved interferon (IFN) response gene signature in inflamed mucosa of children and adults with EoE.(48) While it is tempting to conclude that this finding is a result of IFNγ producing T cells in our culture system,(48) it is notable that the IFN signature was specifically enriched in milk-expanded cells and therefore not a general feature of T cells in culture. An alternative hypothesis is that presence of this IFN response gene signature is indicative of these T cells having circulated through the inflamed esophageal mucosa.

We also observe specific upregulation of genes related to activation (*TMSB4X*, *S100A4*), cytotoxicity (*SRGN*), exhaustion (*CAPG*, *S100A10*), and regulation (*NDFIP1*) in eoeTCR-4 clones compared to the non-milk-specific clones eoeTCR-6 and 7. Of these, S100A4 expression may be of particular relevance as a means to identify food-specific T cell clones, as it has been shown to promote allergic T cell responses and be elevated in EoE.(49,50) Similarly, Ndfip1 is induced when T cells respond to innocuous foreign or self-antigen that should normally be tolerated.(51) Ndfip1 has also been shown to suppress IL-4 production by TH2 cells, and to force their exit from the cell cycle.(52) Ndfip1 may thereby limit effector functions and contribute to T cell anergy (which is potentially consistent with the concurrent upregulation of *CAPG* and *S100A10*).(51) It is reasonable to hypothesize that this constellation of findings is consistent with a (failed) attempt at promoting tolerance by limiting peTH2 expansion and effector function. Whether these findings are representative of *in vivo* biology remains to be established.

Our findings also have broad implications for the development of antigen-specific next-generation diagnostics, biomarkers, and therapeutics for EoE. Though efforts are underway to develop and implement functional T cell assays to detect food allergen-activated T cell responses in the circulation of EoE patients,(11,12) clinical diagnostics to aid in the identification of EoE-causal foods are not currently available. Understanding the TCR repertoire of food antigen recognition in EoE holds the potential to facilitate the development of tetramer-based assays that simultaneously and directly detect the presence of pathogenic T cell populations for multiple foods.(53) In addition to aiding the development of personalized elimination diets, the ability to detect and monitor allergen-specific T cell responses may be a useful prognostic tool to guide risk assessments and help with timing of food allergen reintroduction.

A natural extension of our study will be to determine whether different T cell functional profiles (pathogenic effector vs regulatory) have distinct TCR repertoires and antigen epitope specificities—information directly relevant to future applications of oral immunotherapy designed to impair pathogenic and/or boost regulatory immune responses. Further, knowing the molecular details of antigen-specificity in food allergy is the first step towards the development of cell-specific therapeutics designed to delete peTH2 cells while leaving food-specific regulatory T cells and unrelated TH cells intact.

There are inherent limitations to our study. As our study focuses on the molecular basis of EoE milk allergy, future studies are needed to understand the molecules relevant to the presentation and recognition of other EoE food allergens. Further, our TCR discovery cohort included one EoE milk allergy patient with a “classic” clinical profile. Analysis of EoE milk allergy patients with other clinical profiles (e.g. EoE after oral immunotherapy and patients with resolving/prior EoE) is therefore necessary. In addition, we tested our milk-expanded TCR clonotypes in a control donor who was matched for HLA-DR and DQ, but mismatched for DP. As such, the “negative” clones reported here could be milk-specific but recognize their cognate antigen in the context of HLA-DP. Finally, as we were unable to synthesize the N-terminus of β-casein, there could be additional cognate antigens for eoeTCR-4 that we didn’t identify here.

In summary, we provide the first report of a molecular basis for food allergen presentation and recognition in EoE. While not designed to provide an exhaustive evaluation of the EoE milk allergy TCR repertoire, our study is an important “proof of concept” that will lay the groundwork for future studies. It is known from prior studies of antigen recognition in food allergy that there is disproportionate expansion of the private TCR repertoire in highly allergic individuals.(54) As such, it is perhaps not surprising that we did not detect eoeTCR-4 in our validation cohorts. Nevertheless, our findings do not preclude the existence of a set of public TCRs that are common to EoE milk allergy, and our platform is well suited to more comprehensive investigations of the EoE TCR repertoire.

## Funding

American Partnership for Eosinophilic Disorders and American Academy of Allergy, Asthma, and Immunology HOPE Grant (DAH)

American Partnership for Eosinophilic Disorders Pilot Grant (DAH)

Food Allergy Fund (DAH)

Hartwell Foundation Individual Biomedical Award (DAH)

American Academy of Allergy, Asthma, and Immunology Faculty Development Award (DAH)

Institutional Development Funds from the Children’s Hospital of Philadelphia (DAH)

Funds from Ira & Diana Riklis and Andrew & Talia Day (JMS)

The Yehudai Family Foundation (JMS)

Food Allergy Science Initiative (WGS)

Institutional Development Funds from the Children’s Hospital of Philadelphia to the Center for Applied Genomics (HH)

The Children’s Hospital of Philadelphia Endowed Chair in Genomic Research (HH)

## Author contributions

Conceptualization: DAH

Investigation: JD, AH, HQ, KEC

Formal analysis: JD, AH, HQ, RLC, KEC, WGS, KC, DAH

Data Curation: JD, AH, HQ, KEC

Writing – original draft: DAH

Writing – review and editing: JD, RLC, SJM, WGS, HH, JMS, KC, DAH

Project administration: DAH

Methodology: SJM, WGS, HH, KC, DAH

Resources: WGS, HH, JMS, KC

Visualization: JC, AH, KEC, WGS, HH, KC, DAH

Supervision: WGS, HH, KC, DAH

Funding acquisition: WGS, HH, JMS, KC, DAH

Approval of Final Manuscript: All

## Conflict of interest disclosure statement

JD, JMS, and DAH hold a patent on the use of functional T cell assays for the identification of EoE-causal foods. The remaining authors have no conflicts of interest to disclose.

## ABBREVIATIONS

(EoE): Eosinophilic esophagitis
(T2): Type 2
(TH2): T helper type 2
(scRNA-seq): single cell RNA sequencing
(IRB): Institutional Review Board
(PBMCs): Peripheral blood mononuclear cells
(CFSE): Carboxyfluorescein succinimidyl ester
(TCR): T cell receptor
(HLA): human leukocyte antigen
(scTCR-seq): Paired *TRA* and *TRB* sequencing
(peTH2): pathogenic effector TH2 cells
(IFN): interferon

## ACKNOWLEDGEMENTS

We acknowledge the patients and families who donated their time and biologic samples to allow this work to be possible. We acknowledge the CHOP Center for Applied Genomics for library generation and sequencing. We thank Max Eldabbas, Miranda Wang, and Emileigh Maddox of the Penn Human Immunology Core for assistance with purification of PBMCs (NIH P30 AI045008 and P30 CA016520, RRID:SCR 022380). We thank Erik Nash and the CHOP Proteomics Core Facility (RRID:SCR_023099) for assistance with proteomic analyses. We thank the CHOP Vector Core for assistance with vector generation. We thank Control 1 for two years of regular blood donations which made it possible to optimize our transfection protocol and test several TCRs under many conditions.

## REFERENCES

1. Landres RT, Kuster GG, Strum WB. Eosinophilic esophagitis in a patient with vigorous achalasia. Gastroenterology. 1978 Jun;74(6):1298–301.

2. Straumann A, Spichtin HP, Bernoulli R, Loosli J, Vögtlin J. Idiopathic Eosinophilic Esophagitis: A Frequently Overlooked Disease with Typical Clinical Aspects and Discrete Endoscopic Findings. Schweiz Med Wochenschr. 1994 Aug 20;124(33):1419–29.

3. Furuta GT, Katzka DA. Eosinophilic Esophagitis. N Engl J Med. 2015 Oct 22;373(17):1640–8.

4. Münch R, Kuhlmann U, Makek M, Ammann R, Siegenthaler W. [Eosinophilic esophagitis, a rare manifestation of eosinophilic gastroenteritis]. Schweiz Med Wochenschr. 1982 May 15;112(20):731–4.

5. Chang JW, Kliewer K, Haller E, Lynett A, Doerfler B, Katzka DA, et al. Development of a Practical Guide to Implement and Monitor Diet Therapy for Eosinophilic Esophagitis. Clin Gastroenterol Hepatol. 2023 Jul;21(7):1690–8.

6. Nicholson AG, Li D, Pastorino U, Goldstraw P, Jeffery PK. Full thickness eosinophilia in oesophageal leiomyomatosis and idiopathic eosinophilic oesophagitis. A common allergic inflammatory profile? J Pathol. 1997 Oct;183(2):233–6.

7. Straumann A, Bauer M, Fischer B, Blaser K, Simon HU. Idiopathic eosinophilic esophagitis is associated with a T(H)2-type allergic inflammatory response. JAllergy ClinImmunol. 2001;108(6):954–61.

8. Mishra A, Schlotman J, Wang M, Rothenberg ME. Critical role for adaptive T cell immunity in experimental eosinophilic esophagitis in mice. J Leukoc Biol. 2007 Apr;81(4):916–24.

9. Noti M, Wojno EDT, Kim BS, Siracusa MC, Giacomin PR, Nair MG, et al. Thymic stromal lymphopoietin-elicited basophil responses promote eosinophilic esophagitis. Nat Med. 2013 Aug;19(8):1005–13.

10. Cianferoni A, Ruffner MA, Guzek R, Guan S, Brown-Whitehorn T, Muir A, et al. Elevated expression of activated TH2 cells and milk-specific TH2 cells in milk-induced eosinophilic esophagitis. Ann Allergy Asthma Immunol. 2018 Feb;120(2):177–183.e2.

11. Dellon ES, Guo R, McGee SJ, Hamilton DK, Nicolai E, Covington J, et al. A Novel Allergen-Specific Immune Signature-Directed Approach to Dietary Elimination in Eosinophilic Esophagitis. Clin Transl Gastroenterol. 2019 Dec;10(12):e00099.

12. Dilollo J, Rodríguez-López EM, Wilkey L, Martin EK, Spergel JM, Hill DA. Peripheral markers of allergen-specific immune activation predict clinical allergy in eosinophilic esophagitis. Allergy. 2021 Nov;76(11):3470–8.

13. Wen T, Aronow BJ, Rochman Y, Rochman M, Kc K, Dexheimer PJ, et al. Single-cell RNA sequencing identifies inflammatory tissue T cells in eosinophilic esophagitis. J Clin Invest. 2019 Apr 8;129(5):2014–28.

14. Morgan DM, Ruiter B, Smith NP, Tu AA, Monian B, Stone BE, et al. Clonally expanded, GPR15-expressing pathogenic effector TH2 cells are associated with eosinophilic esophagitis. Sci Immunol. 2021 Aug 13;6(62):eabi5586.

15. Janarthanam R, Kuang FL, Zalewski A, Amsden K, Wang MY, Ostilla L, et al. Bulk T-cell receptor sequencing confirms clonality in pediatric eosinophilic esophagitis and identifies a food-specific repertoire. Allergy. 2023 Sep;78(9):2487–96.

16. Laky K, Kinard JL, Li JM, Moore IN, Lack J, Fischer ER, et al. Epithelial-intrinsic defects in TGFβR signaling drive local allergic inflammation manifesting as eosinophilic esophagitis. Sci Immunol. 2023 Jan 6;8(79):eabp9940.

17. Buch MH, Vital EM, Emery P. Abatacept in the treatment of rheumatoid arthritis. Arthritis Res Ther. 2008;10 Suppl 1(Suppl 1):S5.

18. Reyes-Castillo Z, Palafox-Sánchez CA, Parra-Rojas I, Martínez-Bonilla GE, del Toro-Arreola S, Ramírez-Dueñas MG, et al. Comparative analysis of autoantibodies targeting peptidylarginine deiminase type 4, mutated citrullinated vimentin and cyclic citrullinated peptides in rheumatoid arthritis: associations with cytokine profiles, clinical and genetic features. Clin Exp Immunol. 2015 Nov;182(2):119–31.

19. Nilsson C, Berthold M, Mascialino B, Orme ME, Sjölander S, Hamilton RG. Accuracy of component-resolved diagnostics in peanut allergy: Systematic literature review and meta-analysis. Pediatr Allergy Immunol. 2020 Apr;31(3):303–14.

20. Barber D, Diaz-Perales A, Escribese MM, Kleine-Tebbe J, Matricardi PM, Ollert M, et al. Molecular allergology and its impact in specific allergy diagnosis and therapy. Allergy. 2021 Dec;76(12):3642–58.

21. Dellon ES, Liacouras CA, Molina-Infante J, Furuta GT, Spergel JM, Zevit N, et al. Updated International Consensus Diagnostic Criteria for Eosinophilic Esophagitis: Proceedings of the AGREE Conference. Gastroenterology. 2018 Oct;155(4):1022–1033.e10.

22. Zheng GXY, Terry JM, Belgrader P, Ryvkin P, Bent ZW, Wilson R, et al. Massively parallel digital transcriptional profiling of single cells. Nat Commun. 2017 Jan 16;8:14049.

23. Lun ATL, McCarthy DJ, Marioni JC. A step-by-step workflow for low-level analysis of single-cell RNA-seq data with Bioconductor. F1000Res. 2016;5:2122.

24. immunarch [Internet]. Available from: 10.5281/zenodo.3367200

25. Trapnell C, Cacchiarelli D, Grimsby J, Pokharel P, Li S, Morse M, et al. The dynamics and regulators of cell fate decisions are revealed by pseudotemporal ordering of single cells. Nat Biotechnol. 2014 Apr;32(4):381–6.

26. Qiu X, Mao Q, Tang Y, Wang L, Chawla R, Pliner HA, et al. Reversed graph embedding resolves complex single-cell trajectories. Nat Methods. 2017 Oct;14(10):979–82.

27. Cao J, Spielmann M, Qiu X, Huang X, Ibrahim DM, Hill AJ, et al. The single-cell transcriptional landscape of mammalian organogenesis. Nature. 2019 Feb;566(7745):496–502.

28. Haghverdi L, Lun ATL, Morgan MD, Marioni JC. Batch effects in single-cell RNA-sequencing data are corrected by matching mutual nearest neighbors. Nat Biotechnol. 2018 Jun;36(5):421–7.

29. Mclnnes, L; Healy, J; Melville, J. Uniform Manifold Approximation and Projection for dimension reduction [Internet]. Available from: https://arxiv.org/abs/1802.03426

30. Traag VA, Waltman L, van Eck NJ. From Louvain to Leiden: guaranteeing well-connected communities. Sci Rep. 2019 Mar 26;9(1):5233.

31. Levine JH, Simonds EF, Bendall SC, Davis KL, Amir el-AD, Tadmor MD, Litvin O, Fienberg HG, Jager A, Zunder ER, Finck R, Gedman AL, Radtke I, Downing JR, Pe’er D, Nolan GP. Data-Driven Phenotypic Dissection of AML Reveals Progenitor-like Cells that Correlate with Prognosis. Cell. 2015 Jul 2;162(1):184–97.

32. Linsley PS, Barahmand-Pour-Whitman F, Balmas E, DeBerg HA, Flynn KJ, Hu AK, et al. Autoreactive T cell receptors with shared germline-like α chains in type 1 diabetes. JCI Insight. 2021 Nov 22;6(22):e151349.

33. Reynisson B, Barra C, Kaabinejadian S, Hildebrand WH, Peters B, Nielsen M. Improved Prediction of MHC II Antigen Presentation through Integration and Motif Deconvolution of Mass Spectrometry MHC Eluted Ligand Data. J Proteome Res. 2020 Jun 5;19(6):2304–15.

34. Minimal, encapsulated proteomic-sample processing applied to copy-number estimation in eukaryotic cells - PubMed [Internet]. [cited 2024 Mar 30]. Available from: https://pubmed.ncbi.nlm.nih.gov/24487582/

35. The MaxQuant computational platform for mass spectrometry-based shotgun proteomics - PubMed [Internet]. [cited 2024 Mar 30]. Available from: https://pubmed.ncbi.nlm.nih.gov/27809316/

36. Proteomics Quality Control: Quality Control Software for MaxQuant Results - PubMed [Internet]. [cited 2024 Mar 30]. Available from: https://pubmed.ncbi.nlm.nih.gov/26653327/

37. Non-parametric estimation of posterior error probabilities associated with peptides identified by tandem mass spectrometry - PubMed [Internet]. [cited 2024 Mar 30]. Available from: https://pubmed.ncbi.nlm.nih.gov/18689838/

38. A statistical model for identifying proteins by tandem mass spectrometry - PubMed [Internet]. [cited 2024 Mar 30]. Available from: https://pubmed.ncbi.nlm.nih.gov/14632076/

39. Bolotin DA, Shugay M, Mamedov IZ, Putintseva EV, Turchaninova MA, Zvyagin IV, et al. MiTCR: software for T-cell receptor sequencing data analysis. Nat Methods. 2013 Sep;10(9):813–4.

40. Van Der Loo M. The stringdist Package for Approximate String Matching. The R Journal. 2014 Apr 27;6(1):111–22.

41. Mayer-Blackwell K, Schattgen S, Cohen-Lavi L, Crawford JC, Souquette A, Gaevert JA, et al. TCR meta-clonotypes for biomarker discovery with tcrdist3 enabled identification of public, HLA-restricted clusters of SARS-CoV-2 TCRs. Elife. 2021 Nov 30;10:e68605.

42. Rothenberg ME, Spergel JM, Sherrill JD, Annaiah K, Martin LJ, Cianferoni A, et al. Common variants at 5q22 associate with pediatric eosinophilic esophagitis. NatGenet. 2010;42(4):289–91.

43. Sleiman PM, Wang ML, Cianferoni A, Aceves S, Gonsalves N, Nadeau K, et al. GWAS identifies four novel eosinophilic esophagitis loci. NatCommun. 2014;5(Journal Article):5593.

44. Chang X, March M, Mentch F, Nguyen K, Glessner J, Qu H, et al. A genome-wide association meta-analysis identifies new eosinophilic esophagitis loci. J Allergy Clin Immunol. 2022 Mar;149(3):988–98.

45. Kelly KJ, Lazenby AJ, Rowe PC, Yardley JH, Perman JA, Sampson HA. Eosinophilic esophagitis attributed to gastroesophageal reflux: improvement with an amino acid-based formula. Gastroenterology. 1995;109(5):1503–12.

46. Peterson KA, Byrne KR, Vinson LA, Ying J, Boynton KK, Fang JC, et al. Elemental diet induces histologic response in adult eosinophilic esophagitis. AmJGastroenterol. 2013;108(5):759–66.

47. Kliewer KL, Gonsalves N, Dellon ES, Katzka DA, Abonia JP, Aceves SS, et al. One-food versus six-food elimination diet therapy for the treatment of eosinophilic oesophagitis: a multicentre, randomised, open-label trial. Lancet Gastroenterol Hepatol. 2023 May;8(5):408–21.

48. Ruffner MA, Hu A, Dilollo J, Benocek K, Shows D, Gluck M, et al. Conserved IFN Signature between Adult and Pediatric Eosinophilic Esophagitis. J Immunol. 2021 Mar 15;206(6):1361–71.

49. Bruhn S, Fang Y, Barrenäs F, Gustafsson M, Zhang H, Konstantinell A, et al. A generally applicable translational strategy identifies S100A4 as a candidate gene in allergy. Sci Transl Med. 2014 Jan 8;6(218):218ra4.

50. Babble J, Dong S, Lewis N, Mo J, Faulkner C, Chiang A, et al. S100A4 Levels in Pediatric Eosinophilic Esophagitis Cohort. Journal of Allergy and Clinical Immunology. 149(2):AB157.

51. Altin JA, Daley SR, Howitt J, Rickards HJ, Batkin AK, Horikawa K, et al. Ndfip1 mediates peripheral tolerance to self and exogenous antigen by inducing cell cycle exit in responding CD4+ T cells. Proc Natl Acad Sci U S A. 2014 Feb 11;111(6):2067– 74.

52. Oliver PM, Cao X, Worthen GS, Shi P, Briones N, MacLeod M, et al. Ndfip1 protein promotes the function of itch ubiquitin ligase to prevent T cell activation and T helper 2 cell-mediated inflammation. Immunity. 2006 Dec;25(6):929–40.

53. Sarna VK, Lundin KEA, Mørkrid L, Qiao SW, Sollid LM, Christophersen A. HLA-DQ-Gluten Tetramer Blood Test Accurately Identifies Patients With and Without Celiac Disease in Absence of Gluten Consumption. Gastroenterology. 2018 Mar;154(4):886–896.e6.

54. Ruiter B, Smith NP, Monian B, Tu AA, Fleming E, Virkud YV, et al. Expansion of the CD4+ effector T-cell repertoire characterizes peanut-allergic patients with heightened clinical sensitivity. J Allergy Clin Immunol. 2020 Jan;145(1):270–82.

