## Extended Data Tables for "Companion to: A molecular basis for milk allergen immune recognition in eosinophilic esophagitis"

### Extended Data Table 1 | TRA and TRB gene usage for the top 20 milk-expanded clonotypes

| TRA junction | TRAV gene | TRAJ gene | TRB junction | TRBV gene | TRBJ gene | Milk stimulation count | Unstimulated count |
|---|---|---|---|---|---|---|---|
| CALKGTNAGKSTF | TRAV16 | TRAJ27 | CSDKTLHF; CASSLARGSLNTEAFF | TRBV29-1; TRBV7-9 | TRBJ1-6; TRBJ1-1 | 13 | 8 |
| CAVEDRGGATNKLIF; CVVVGRTGGFKTIF | TRAV2; TRAV12-1 | TRAJ32; TRAJ9 | CASSLARGRTDTQYF | TRBV27 | TRBJ2-3 | 14 | 2 |
| CAATFSGSARQLTF | TRAV13-1 | TRAJ22 | CASSQESGAGNTIYF | TRBV4-1 | TRBJ1-3 | 19 | 2 |
| CVVTANSGGSNYKLTF | TRAV8-2 | TRAJ53 | CASSPGLAGAEQYF | TRBV5-4 | TRBJ2-7 | 7 | 5 |
| CALSDQGAQKLVF | TRAV9-2 | TRAJ54 | CASSWVLGGSYEQYF | TRBV27 | TRBJ2-7 | 12 | 0 |
| CAFSNNDMRF | TRAV38-1 | TRAJ43 | CSASQGGGQNYGYTF | TRBV20-1 | TRBJ1-2 | 7 | 3 |
| CAFMNIGSNTGKLIF | TRAV38-1 | TRAJ37 | CASSGQGYEQYF | TRBV6-6 | TRBJ2-7 | 10 | 0 |
| CAPFSGYALNF | TRAV21 | TRAJ41 | CASSPGQLFTYEQYF | TRBV11-2 | TRBJ2-7 | 8 | 1 |
| CALKNNNARLMF | TRAV16 | TRAJ31 | CASSLGLAGVGLSYEQYF | TRBV5-1 | TRBJ2-7 | 6 | 3 |
| CGTEKGSGSRLTF | TRAV30 | TRAJ58 | CASSPSIRYEQYF | TRBV6-5 | TRBJ2-7 | 8 | 0 |
| CATDAGAQKLVF | TRAV17 | TRAJ54 | CASSPARTGGGYGYTF | TRBV5-5 | TRBJ1-2 | 8 | 0 |
| CALSVRGFGNVLHC | TRAV19 | TRAJ35 | CASSSDTRGGTQYF | TRBV27 | TRBJ2-3 | 7 | 1 |
| CAIHDTASKLTF | TRAV17 | TRAJ44 | CASSTWGASSYEQYF | TRBV6-5 | TRBJ2-7 | 8 | 0 |
| CAEKSSGNQFYF | TRAV13-2 | TRAJ49 | CASRKWGESEKLFF | TRBV27 | TRBJ1-4 | 8 | 0 |
| CAAAQNDYKLSF | TRAV17 | TRAJ20 | CASSISGSYEQYF | TRBV19 | TRBJ2-7 | 8 | 0 |
| CAVPSHQGGKLIF | TRAV2 | TRAJ23 | CASSQPSGGAGETQYF | TRBV3-1 | TRBJ2-5 | 7 | 0 |
| CAGPGAGNNRKLIW | TRAV35 | TRAJ38 | CSANGSPTKGEQYF | TRBV20-1 | TRBJ2-7 | 7 | 0 |
| CILKSSYYIQGAQKLVF | TRAV26-2 | TRAJ54 | CASSVYSSEVDTQYF | TRBV9 | TRBJ2-3 | 6 | 0 |
| CAVNAGAAGNKLTF | TRAV8-1 | TRAJ17 | CASSESGTDRAMFF | TRBV10-1 | TRBJ2-2 | 6 | 0 |
| CAVGAPGGSYIPTH | TRAV8-3 | TRAJ6 | CAWSVGENQPQHF | TRBV30 | TRBJ1-5 | 6 | 0 |

### Extended Data Table 2 | Demographics and HLA Genotyping of subjects

| Subject | Age | Sex | Race | Ethnicity | DRB1 | DR additional | DPA1 | DPB1 | DQA1 | DQB1 | A | B | C |
|---|---|---|---|---|---|---|---|---|---|---|---|---|---|
| Control 1 | 30 | M | White | NH/L | 07:01, 11:01 | DRB3 02:02, DRB4 01:01 | 01:03, 01:03 | 03:01, 04:01 | 02:01, 05:05 | 02:02, 03:01 | 01:01, 29:02 | 13:02, 44:03 | 06:02, 16:01 |
| Control 2 | 40 | M | White | NH/L | 07:01, 13:01 | DRB3 02:02, DRB4 01:03 | 01:03, 01:03 | 04:01, 04:02 | 01:03, 02:01 | 03:03, 06:03 | 01:01, 02:01 | 56:01, 57:01 | 01:02, 06:02 |
| Control 3 | 28 | F | ND | ND | 11:03, 15:01 | DRB3 02:02 | 01:03, 01:03 | 04:01, 04:01 | 01:02, 05:05 | 03:01, 06:02 | 03:01, 24:02 | 07:02, 15:01 | 03:03, 07:02 |
| Control 4 | 29 | M | ND | ND | 04:01, 04:03 | DRB4 01:03 | 02:02, 02:02 | 05:01, 05:01 | 03:01, 03:01 | 03:02, 03:02 | 03:01, 24:02 | 40:01, 52:01 | 03:03, 12:02 |
| EoE 1 | 14 | M | White | NH/L | 07:01, 11:04 | DRB3 02:02, DRB4 01:03 | 01:03, 01:04 | 02:01, 15:01 | 02:01, 05:05 | 03:01, 03:03 | 01:01, 03:01 | 07:02, 57:01 | 07:02, 06:02 |

NH/L = Non-Hispanic or Latino; H/L = Hispanic or Latino; ND = No data; N/A = Not applicable

### Extended Data Table 3 | β-casein peptide library and reactivity with eoeTCR-4

| Peptide | Amino acids | Sequence | Length | Notes | Reactivity with eoeTCR-4 |
|---|---|---|---|---|---|
| | 1-56 | MKVLILACLVALALARELEELNVPGEIV ESLSSSEESITRINKKIEKFQSEEQQQT | 56 AA | Unable to synthesize | Unknown |
| 1 | 59-78 | ELQDKIHPFAQTQSLVYPFP | 20 AA | Contain DRB1*07:01 predicted bind site FAQTQSLVY | Yes |
| 2 | 76-95 | PFPGPIPNSLPQNIPPLTQT | 20 AA | | No |
| 3 | 93-112 | TQTPVVVPPFLQPEVMGVSK | 20 AA | | No |
| 4 | 110-129 | VSKVKEAMAPKHKEMPFPKY | 20 AA | | No |
| 5 | 127-143 | PKYPVEPFTESQSLTLT | 17 AA | Contain DRB1*07:01 predicted bind site FTESQSLTL | No |
| 6 | 130-150 | PVEPFTESQSLTLTDVENLHL | 20 AA | Contain DRB1*07:01 predicted bind site FTESQSLTL | No |
| 7 | 140-160 | LTLTDVENLHLPLPLLQSWMH | 20 AA | | No |
| 8 | 150-170 | LPLPLLQSWMHQPHQPLPPTV | 20 AA | | No |
| 9 | 169-224 | TVMFPPQSVLSLSQSKVLPVPQKAVPYP QRDMPIQAFLLYQEPVLGPVRGPFPIIV | 56 AA | Contain DRB1*07:01 predicted bind site LSQSKVLPV | No |

AA = Amino acids

### Extended Data Table 4 | Demographic characteristics of WES cohort

| Subjects | Birth Year [Median (ranges)] | Sex | Race | Ethnicity |
|---|---|---|---|---|
| EoE (n=132) | 2003 (1992, 2016) | Male 110, Female 22 | White 96, Black 27, Other 9 | Non-Hispanic 118, Hispanic 10, Other 4 |
