## Extended Data Figures for "Companion to: A molecular basis for milk allergen immune recognition in eosinophilic esophagitis"

### Dilollo et al - Extended Data Figure 1

**a** EoE Subject 1 demographic and clinical characteristicts

| Subject | Age at sampling (years) | Sex | Race | Ethnicity | EoE causal foods | Eos/HPF at diagnosis | Eos/HPF at sampling | Other allergic Dx |
|---|---|---|---|---|---|---|---|---|
| EoE 1 | 14 | M | White | Non-Hispanic or Latino | Milk | 20 | 30 | Asthma, AR, IgE-FA, AD |

**b** EoE Subject 1 clinical timeline

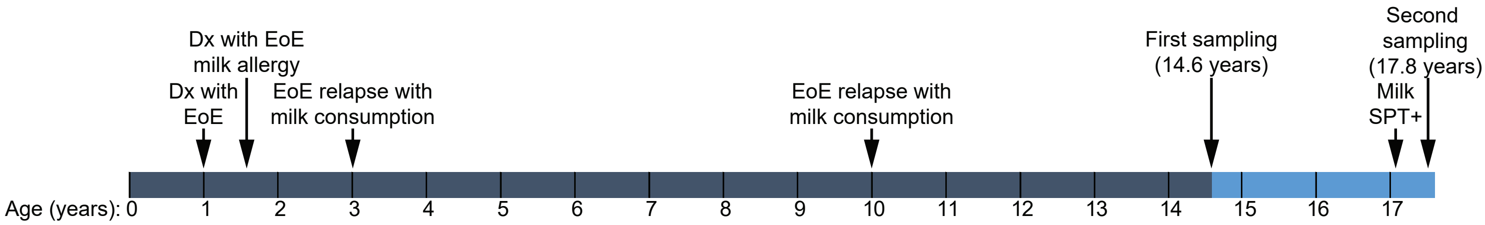


**Extended Data Figure 1. EoE Subject 1 demographic and clinical characteristicts with timeline.** (**a**) EoE Subject 1 demographic and clinical characteristicts. (**b**) EoE Subject 1 clinical timeline. SPT = Skin prick test.

### Dilollo et al - Extended Data Figure 2

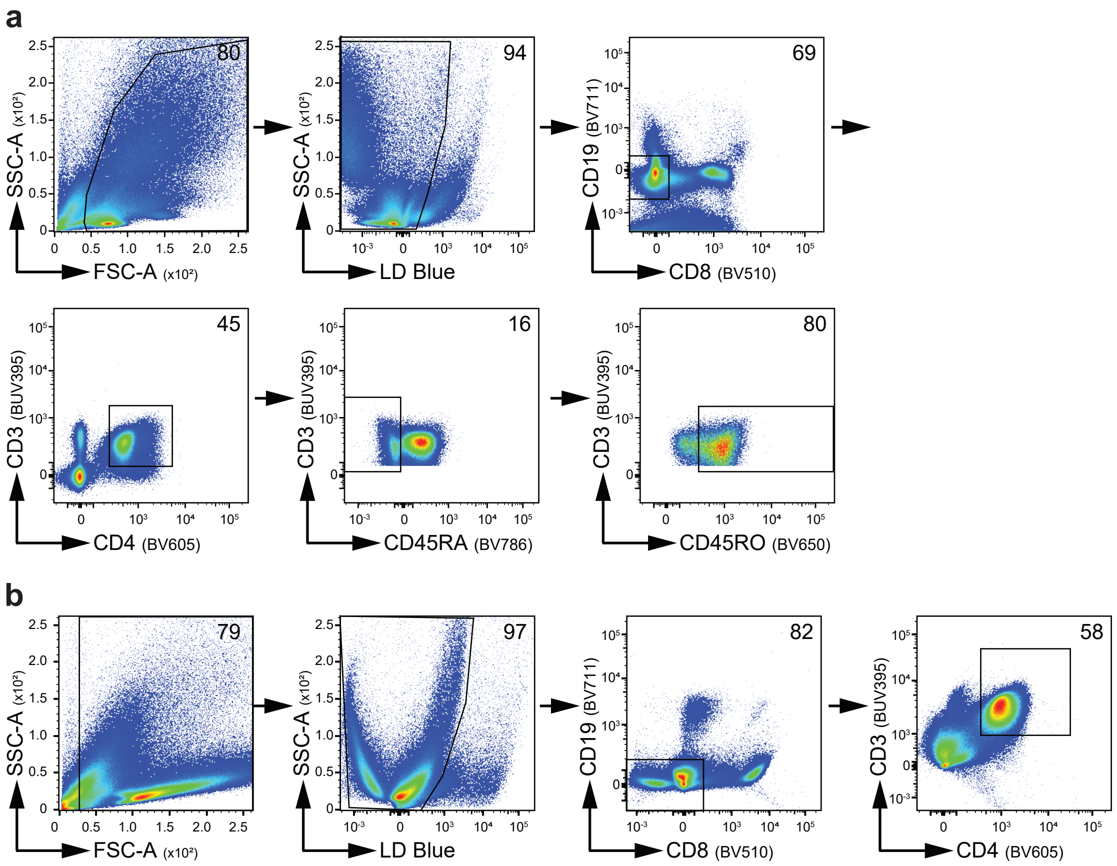


**Extended Data Figure 2. Flow cytometry gating strategies.** (**a**) Gating strategy for PBMC cultures. (**b**) Gating strategy for T cell transduction cultures.

### Dilollo et al - Extended Data Figure 3

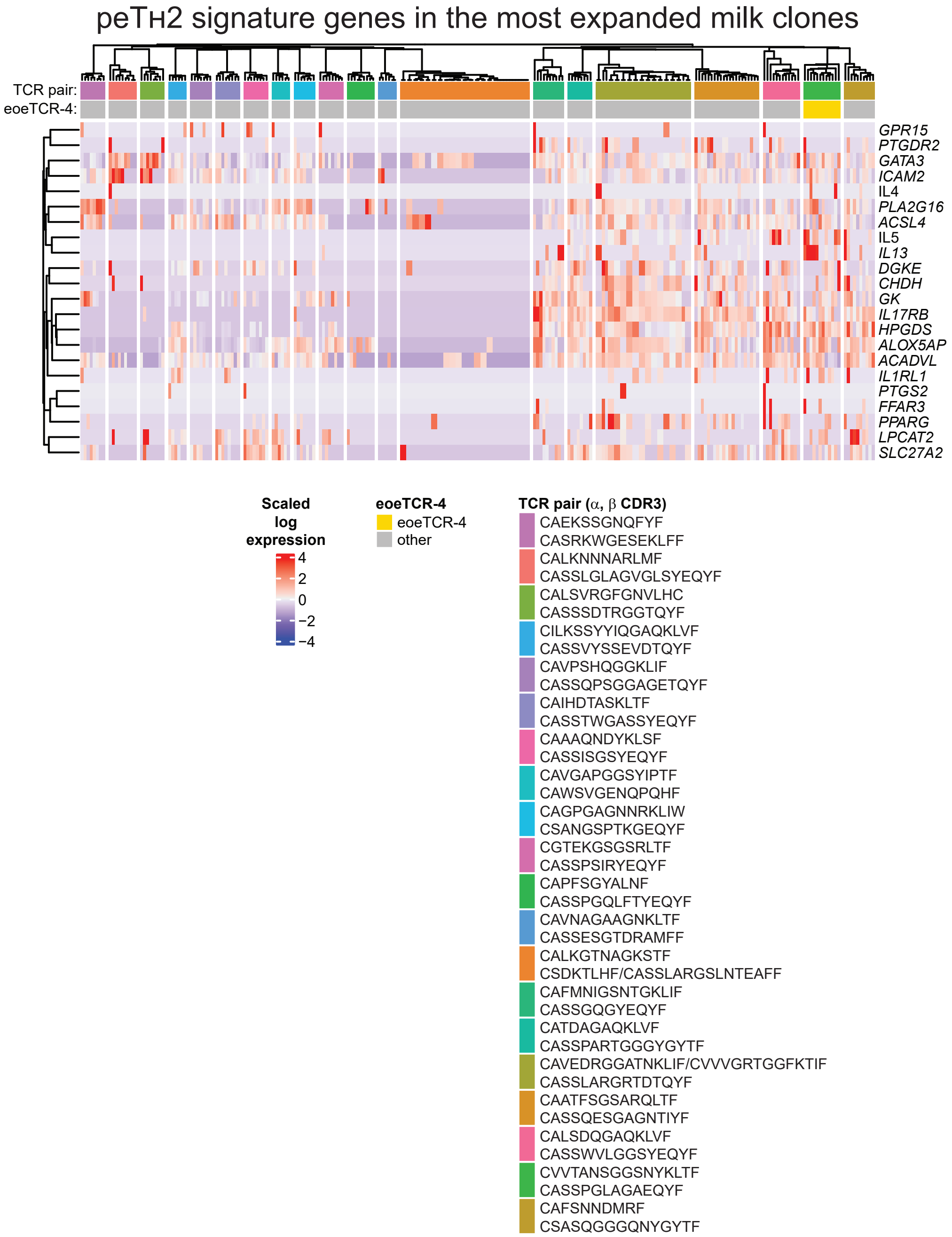


**Extended Data Figure 3. Type 2 gene expression in the 20 most expanded clones from the milk-stimulated CD4+ memory T cell cultures.** peTH2 signature genes in the top 20 most expanded CD4+ memory T cells clonotypes from the milk-stimulated cultures. Scaled log expression, FDR ≤ 0.05.

### Dilollo et al - Extended Data Figure 4

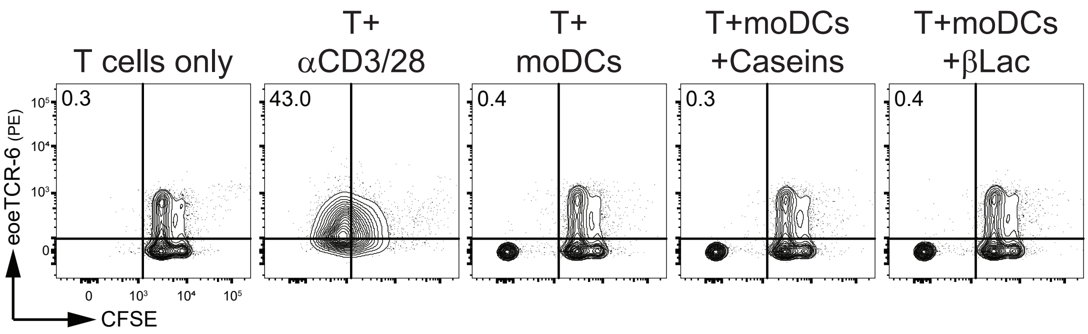


**Extended Data Figure 4. Evaluation of eoeTCR-6.** Carboxyfluorescein succinimidyl ester (CFSE) fluorescence intensity of HLA-DR-matched conventional T cells (T) transduced with eoeTCR-6 and cultured alone or in the presence of anti-CD3 and anti-CD28 beads (αCD3/28), autologous monocyte-derived dendtritic cells (moDCs), or moDCs pulsed with a cocktail of α–casein, β–casein, κ–casein (Caseins) or β–lactoglobulin (βLac).

### Dilollo et al - Extended Data Figure 5

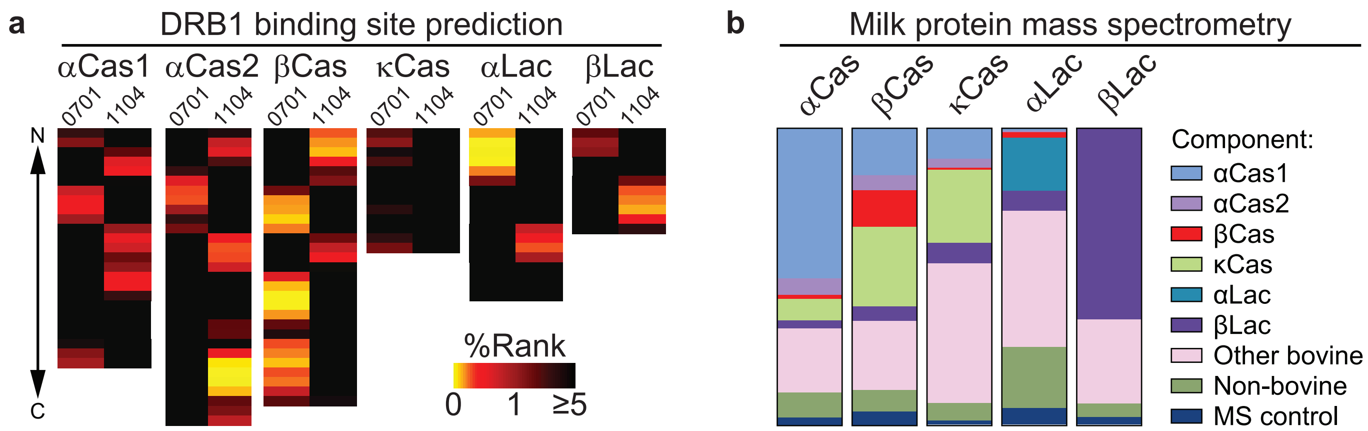


**Extended Data Figure 5. Milk antigen prediction and mass spectrometry.** (**a**) DRB1 binding site prediction for α–casein1 (αCas1), α–casein2 (αCas2), β–casein (βCas), κ–casein (κCas), α–lactalbumin (αLac), or β–lactoglobulin (βLac) by HLA-DR allele (% Rank score normalized to a set of randomly generated peptides, 0-1 “strong bind”, 1-5 “weak bind”, >5 “no bind” as determined by NetMHCIIpan). (**b**) Mass spectrometry of purified milk protein preparations.

### Dilollo et al - Extended Data Figure 6

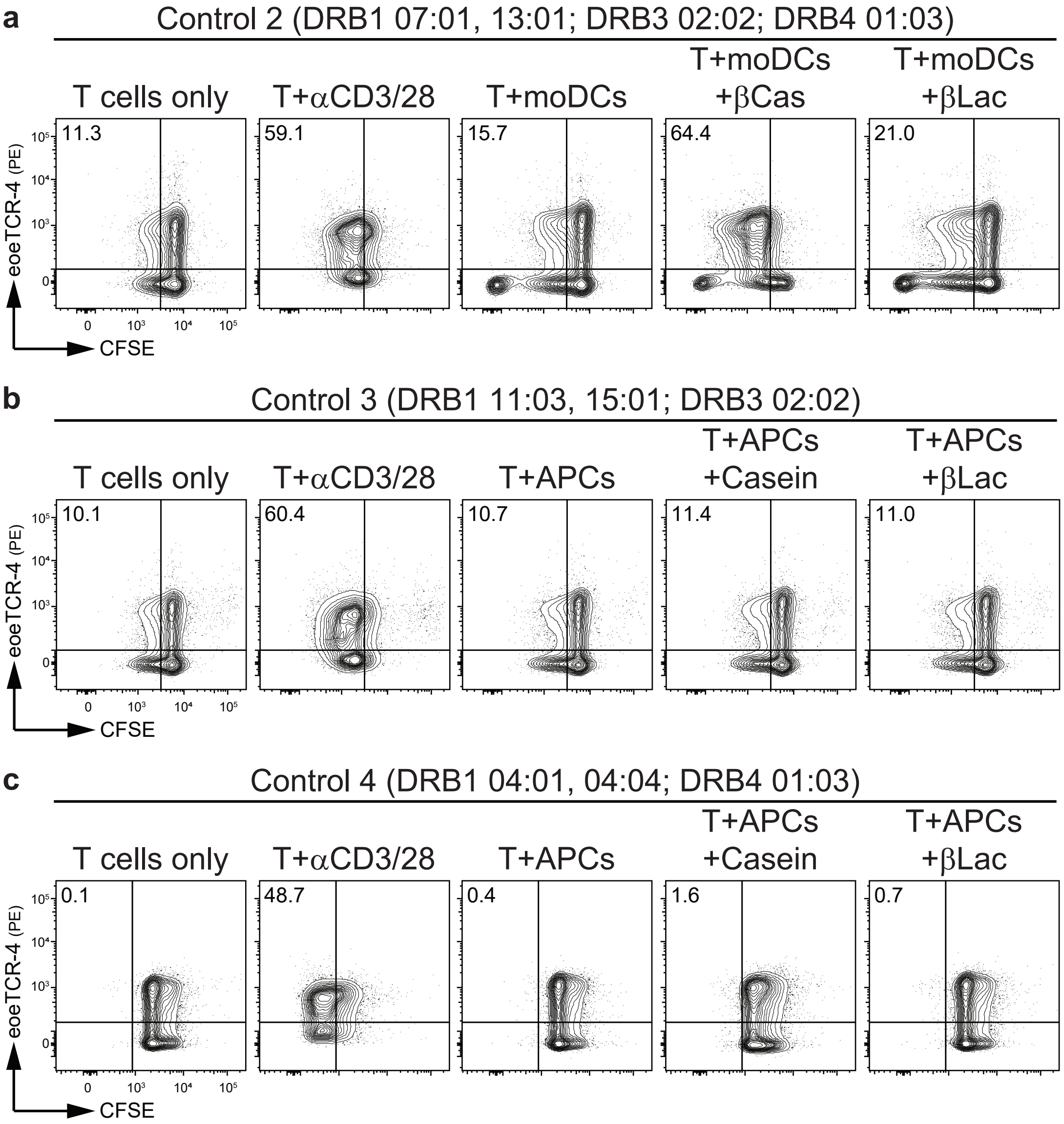


**Extended Data Figure 6. Milk protein presentation via eoeTCR-4 is restricted to HLA-DRB1 07:01.** (**a**) Carboxyfluorescein succinimidyl ester (CFSE) fluorescence intensity of conventional T cells (T) from an individual with the 07:01, 13:01 HLA-DRB1 haplotype transduced with eoeTCR-4 and cultured alone or in the presence of anti-CD3 and anti-CD28 beads (αCD3/28), autologous monocyte-derived dendtritic cells (moDCs), or moDCs pulsed with β–casein (βCas) or β–lactoglobulin (βLac). (**b**) CFSE fluorescence intensity of T cells from an individual with the 11:03, 15:01 HLA-DRB1 haplotype transduced with eoeTCR-4 and cultured alone or in the presence of αCD3/28, autologous antigen presenting cells (APCs), or APCs pulsed with a cocktail of α–casein, β–casein, κ–casein (Casein) or βLac. (**c**) CFSE fluorescence intensity of T cells from an individual with the 04:01, 04:04 HLA-DRB1 haplotype transduced with eoeTCR-4 and cultured alone or in the presence of αCD3/28, APCs, or APCs pulsed with Casein or βLac.

Dilollo et al - Extended Data Figure 7

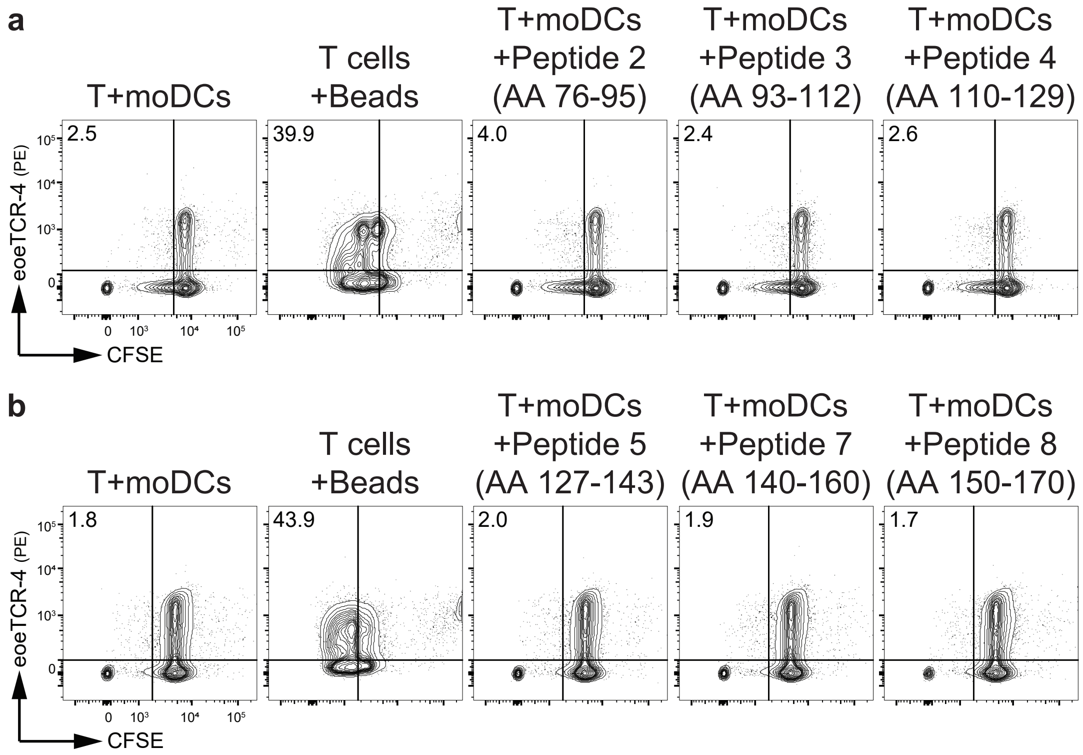


**Extended Data Figure 7. β–casein peptide library screen for eoeTCR-4.** (**a**) Carboxyfluorescein succinimidyl ester (CFSE) fluorescence intensity of T cells transduced with eoeTCR-4 and cultured with moDCs alone, anti-CD3 anti-CD28 activating beads (beads), or moDCs pulsed with β–casein peptides 2, 3, or 4. (**b**) CFSE fluorescence intensity of T cells transduced with eoeTCR-4 and cultured with moDCs alone, beads, or moDCs pulsed with β–casein peptides 5, 7, or 8. Data representative of 2 or more independent experiments.

### Dilollo et al - Extended Data Figure 8

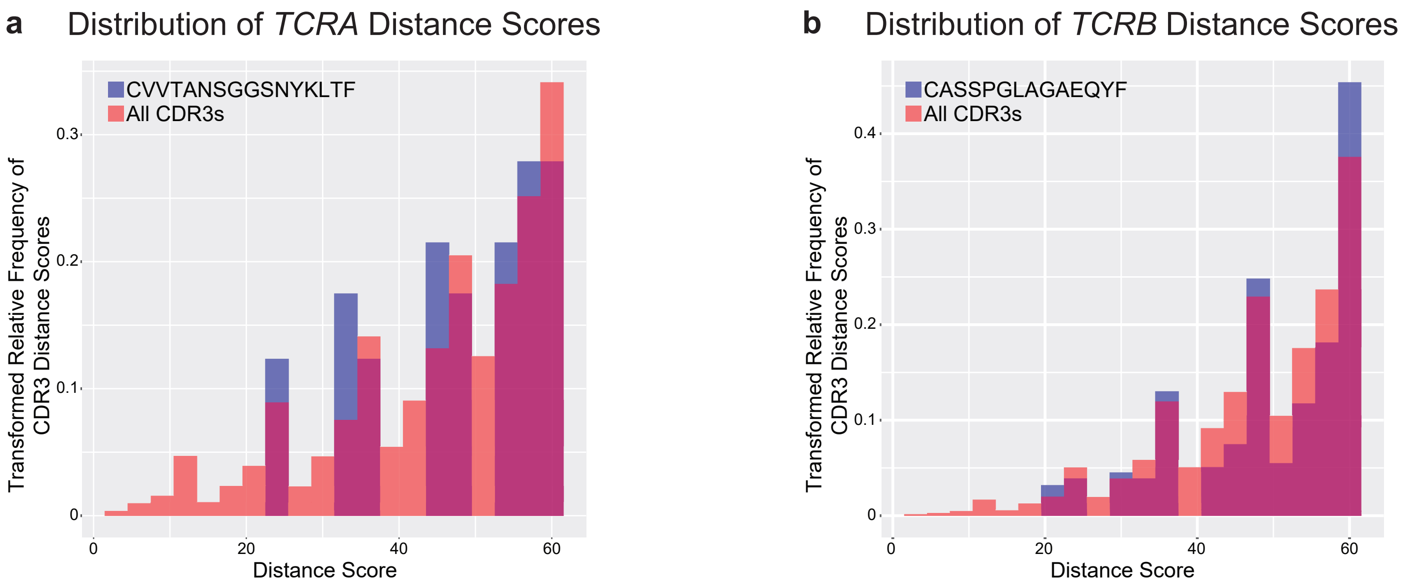


**Extended Data Figure 8. Relative frequency of TCR distance scores.** Histograms illustrate the distance scores (x-axis) computed with tcrdist3 for (**a**) *TCRA* and (**b**) *TCRB* CDR3 sequences against the arcsine square root transformed relative frequency of each score. Orange bars depict transformed distance scores between CDR3s derived from a composite dataset comprising 21 esophageal tissue and PBMC samples obtained from patients with EoE milk allergy. Blue bars depict distance scores between eoeTCR-4 and the composite dataset. Maroon bars depict overlap.

Dilollo et al - Extended Data Figure 9

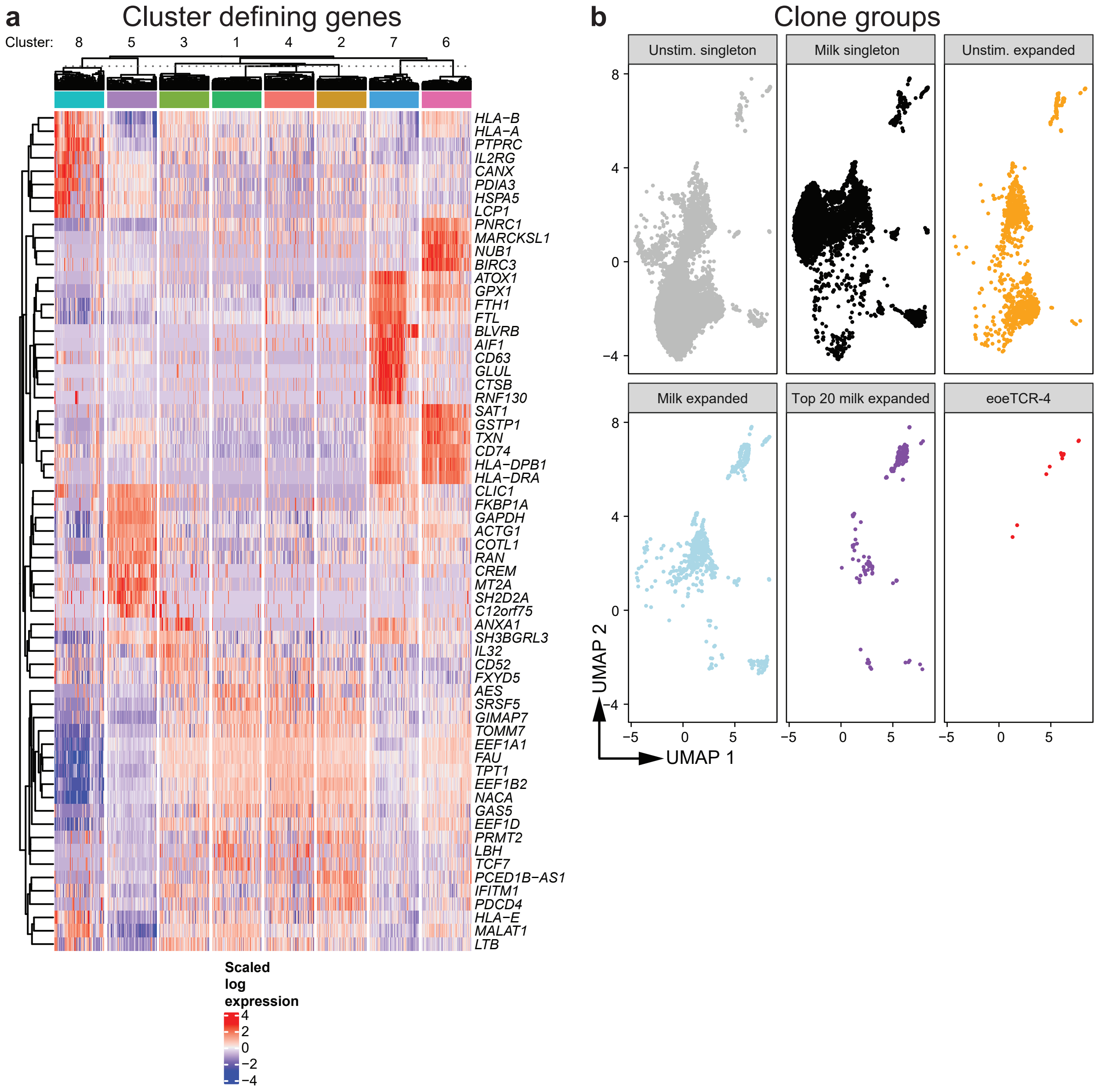


**Extended Data Figure 9. UMAP cluster defining genes and clone group distributions.** (**a**). UMAP cluster defining genes. Scaled log expression, FDR ≤ 0.05. (**b**) Individual UMAPs of CD4+ memory T cell clonotype groups from unstimulated and milk-stimulated culture conditions.

### Dilollo et al - Extended Data Figure 10

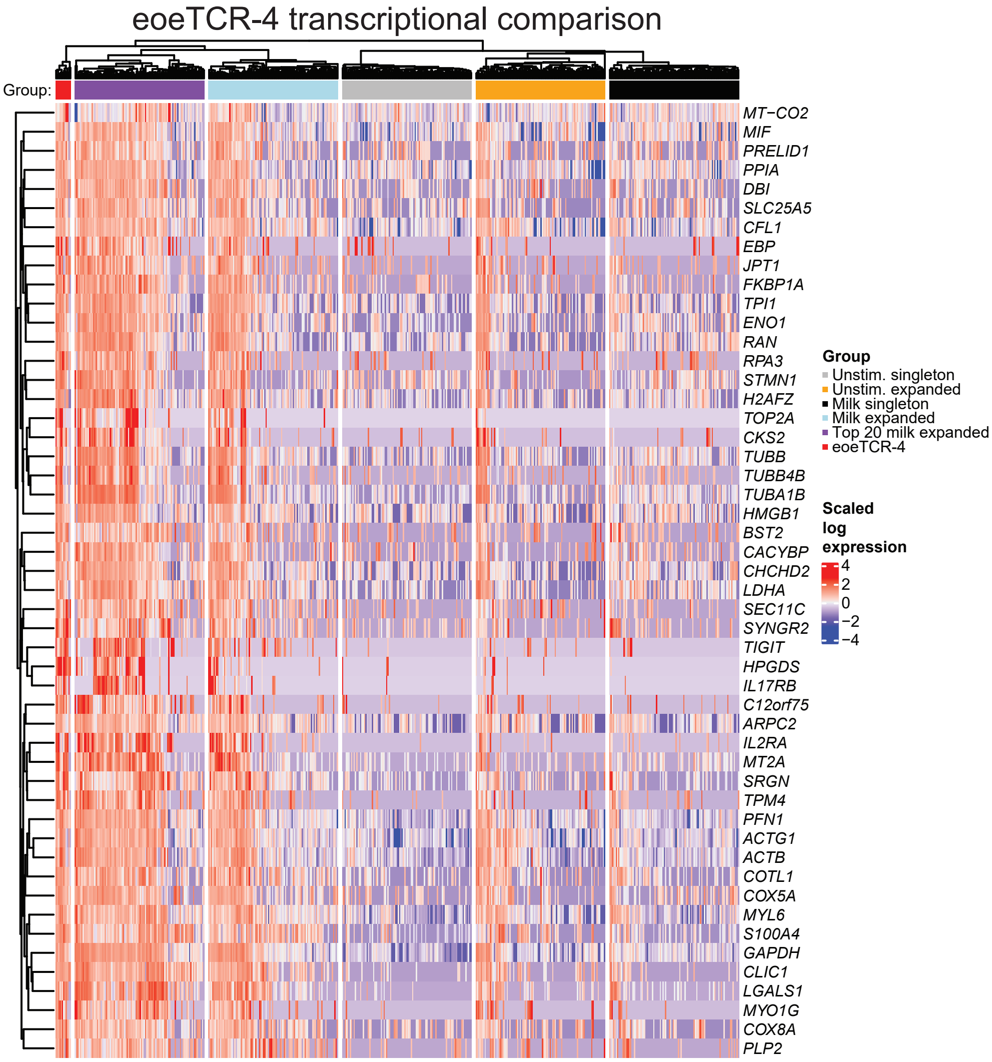


**Extended Data Figure 10. Transcriptional comparison of eoeTCR-4 vs T cell clonotype groups.** Scaled log expression of differentially expressed genes between cells expressing eoeTCR-4 and unstimulated unexpanded cells, displayed in random subsamples of 100 cells per experimental group. FDR ≤ 0.05.
